# *In silico* characterization of unique fungal modular rhodopsin expands the horizon of novel optobiological and biomedical applications

**DOI:** 10.64898/2026.05.25.727616

**Authors:** Alka Kumari, Abhishek Kumar, Komal Sharma, Swaroop Ranjan Pati, Shilpa Mohanty, Suneel Kateriya

## Abstract

Microbial modular rhodopsins, in which light-sensing rhodopsin domains are fused with effector modules, have emerged as promising tools for optogenetic regulation in algae and other systems. However, the diversity and potential regulatory roles of fungal modular rhodopsins remain largely unexplored. Here, we performed a comprehensive *in-silico* analysis to identify previously uncharacterized fungal modular-rhodopsins that pair a conserved light-sensing core with diverse effector domains, including RPEL-motif, NADP-binding Rossmann fold domain, MCM (Mini-Chromosome Maintenance) domain, and GC-cAT (Carnitine O-Acetyltransferase) modules. In *Aureobasidium pullulans*, the representative modular rhodopsin (ApRh-RPEL) contains RPEL-motif associated with actin-related and transcriptional regulatory processes, suggesting light-driven fungal signaling pathway involved in transcriptional and cellular regulation, respectively. Rhodopsins fused with NADP-binding Rossmann fold and MCM domains further indicate possible applications in light-programmable metabolic and cell-cycle signaling. Genome mining additionally revealed that *A. pullulans* harbours a diverse but underexplored array of biosynthetic gene clusters (BGCs), raising the intriguing possibility that light perception may regulate secondary metabolite pathways. Supporting this, multisource protein-protein interaction network analysis links ApRh-RPEL to enzymes involved in terpenoid and sphingolipid biosynthesis, indicating potential cross-talk between light-sensing module and metabolic regulation. These findings outline a computationally derived model in which fungal modular rhodopsins (ApRh-RPEL) function as opto-synthetic regulators of biosynthetic processes. Structural predictions confirmed conserved Schiff-base lysine and retinal-binding pocket, highlighting functional diversity across fungal rhodopsins. Together, these findings expand the optogenetic toolkit and provide a framework for engineering light-driven signaling in fungi, with applications in optobiological and biomedical applications.

## Introduction

Light is a fundamental energy source for ecosystems, driving photosynthesis and supporting the survival of non-photosynthetic organisms by acting as an environmental cue detected through photoreceptors (Rao and Xue 2024). Among these, rhodopsins are ubiquitous heptahelical transmembrane photoreceptors. They are classified into two families: microbial (type I) and animal (type II), both of which share a conserved seven-transmembrane architecture but differ in their retinal configurations and signaling mechanisms. Microbial rhodopsins typically employ all-*trans*/13-*cis* retinal and function as light-driven ion pumps, channels, or sensory receptors, exemplified by bacteriorhodopsin (BR) from *Halobacterium salinarum*. In contrast, type II rhodopsins are GPCRs that use 11-*cis*/all-*trans* retinal to initiate visual phototransduction. The discovery of channelrhodopsin (ChR) from *Chlamydomonas reinhardtii* expanded interest in microbial rhodopsins, highlighting their versatility as light-regulated ion transporters and signaling modules (Emiliani et al. 2022; Ernst et al. 2014; Kulbay et al. 2024; Liu and Colmenares 2003; Nagata and Inoue 2021). The retinal-dependent photochemistry provides high spatiotemporal resolution, low toxicity, and tunable wavelength responsiveness, making them ideal for neural modulation and emerging photodynamic applications (Vickerman et al. 2021). Beyond classical ion transporters, newer microbial rhodopsins expand this toolkit by mediating flux of diverse cations and anions (Na□, K□, Rb□, Cs□, Cl□) (Inoue et al. 2013; Kandori 2020; Kandori et al. 2018; Kato et al. 2015; Konno et al. 2016; Vickerman et al. 2021), and by functioning as enzyme-coupled photoreceptors such as rhodopsin-guanylyl cyclases (RhGCs) that regulate intracellular second-messenger levels in response to light (Avelar et al. 2014). The continued search for rhodopsins with novel architectures or regulatory modules is therefore essential for advancing, optogenetic, and synthetic-biology applications. Light-sensing mechanisms have independently evolved across archaea, fungi, and animals. In fungi, light plays a central role in regulating development, including conidiation, pigmentation, and circadian rhythms. Fungal photobiology spans a broad spectral range from UV to far-red light, and several rhodopsins have been characterized, including NOP-1 in *Neurospora crassa*, rhodopsins in *Leptosphaeria maculans* and *Allomyces reticulatus*, and rhodopsin–guanylyl cyclases (RhGCs) in zoospore-forming chytrids such as *Blastocladiella emersonii* and *Rhizoclosmatium globosum*, where light-induced cGMP production influences phototactic behaviour (Avelar et al. 2014; Furutani et al. 2004; Purschwitz et al. 2006; Scheib et al. 2015; Waschuk et al. 2005). More recently, a rhodopsin-phosphodiesterase (RhPDE) from the choanoflagellate *Salpingoeca rosetta* was shown to hydrolyze cAMP and cGMP in a light-dependent manner representing the light-activated PDE (Yoshida et al. 2017). These discoveries illustrate the emergence of modular rhodopsins in which light detection is directly coupled to enzymatic or signaling outputs, highlighting the functional diversity of rhodopsin-based photoreception in microorganismsTherefore, it is imperative to analyse the fungal genome to identify novel modular rhodopsins that may mediate varying functions within the cell, apart from maintaining the guanylyl cyclase activity. Hence, the new discovery and identification of novel modular fungal rhodopsins in this study based on bioinformatics tools pave the way for designing new optogenetic tools.

With this objective in view, we examined MycoCosm (fungal genome database), NCBI, and UniProt, leading to the discovery of multiple new modular fungal rhodopsins. We focused on identifying novel modular rhodopsins beyond the already reported RhGC activity. Our analysis unravelled novel modular fungal rhodopsins possessing unique effector domains; these rhodopsin-coupled domains were Rh-RPEL (RPXXXEL motif), Rh-Mini-chromosome Maintenance (MCM)/pyridoxal 5-phosphate (PLP)-dependent enzyme (later referred to as Rh-MCM), Rh-NAD(P) binding Rossmann fold domain, and Rh-guanylyl cyclase coupled with carnitine O-acyltransferase (Rh-GC-cAT). Each of these effector domains was predicted to perform its own unique light-driven functions. The sequence homology and structural homology studies of these proteins showed that they belonged to the Type-I microbial rhodopsins. Interestingly, for the first time, the novel modular domains were reported from the phylum Ascomycota, which have usually been reported in Chytridiomycota. This study has two major highlights. First, a novel *Cladophialophora carrionii* halorhodopsin (ClacRh) and *Boothiomyces macroporosus* channelrhodopsin-like sequences were identified. Previous literature reports have identified only bacteriorhodopsin and sensory rhodopsins in fungi. Second, we identified novel twin rhodopsin-coupled with guanylyl cyclase (RhGC-RhGC) that has not been described before. Further, upon analysis, these rhodopsins were categorized based on their similarities to the well-established class of rhodopsins, including bacteriorhodopsin (BR), *Leptosphaeria* rhodopsin (LR), channelrhodopsin-2 (ChR2), and *Anabaena* sensory rhodopsin (ASR). The *in silico* identification of these novel effector domains in the Kingdom Fungi highlights immense biological, optogenetic, and biotechnological potential. This study lays the foundation for the study of fungal modular rhodopsins that have been demonstrated to modulate the cytoskeletal biology, DNA metabolism, secondary metabolite production and cellular metabolism. Additionally, thorough bioinformatics investigations offer support for the evolutionary diversification and functional adaptation of these newly discovered modular rhodopsins. The presence of these novel fungal modular rhodopsins unravels the unexplored branch of optogenetics that sheds insight into the regulatory role of light in the microbial world. To the best of our knowledge, this is the first *in silico* report of these identified novel modular rhodopsins in fungi.

## Materials and Methods

### Integrative database analysis uncovers novel modular rhodopsin across fungal genomes

The BR and halorhodopsin (HR0 from *Halobacterium salinarum*, ASR (*Nostoc*), and *Chlamydomonas reinhardtii* ChR2 were used as a reference protein and submitted as a query for BLASTp (Altschul et al. 1990) searches in the NCBI (National Centre for Biotechnology Information) non-redundant protein sequence database, JGI MycoCosm database (https://mycocosm.jgi.doe.gov), and Uniprot as of 31^st^ May 2025. The sequences that exhibit significant E-value and maximum coverage were selected for further analysis. The study focused on identifying a new type of modular rhodopsin from fungal sources. Further, the sequences were utilized to identify conserved motifs by comparing the acquired sequences with the NCBI curated database (Wang et al. 2023) using the Conserved Domain Database (CDD) via the CDART (Conserved Domain Architecture Retrieval Tool) (https://www.ncbi.nlm.nih.gov/Structure/lexington/lexington.cgi) to check the authenticity of rhodopsin proteins coupled with diverse effector domains. The determination of the transmembrane helices present within the identified modular rhodopsins was carried out using the transmembrane helices prediction tool, DeepTMHMM-1.0 server (https://services.healthtech.dtu.dk/services/DeepTMHMM-1.0/). Using this tool, the rhodopsin sequence of each identified modular rhodopsin was used as a query sequence in the FASTA format (Hallgren et al. 2022). Based on the results obtained by the server, the schematic representation of the rhodopsin’s transmembrane helices was prepared.

### The first experimental transcript evidence of novel modular rhodopsin from fungal genome database

The transcripts of the putative rhodopsin genes were analyzed using the Joint Genome Institute (JGI) MycoCosm fungal genome portal (https://mycocosm.jgi.doe.gov). The Genome Browser tool was used to visualize transcript evidence, and domain architecture within assembled genomes. Default parameters were applied for BLASTP (BLOSUM62 substitution matrix, word size = 3, gap opening penalty = 11, and extension penalty = 1), with an E-value cutoff of 1 × 10□□ to ensure high-confidence matches. The resulting alignment tables were examined for sequence identity, alignment score, E-value, and subject coverage, with homologous proteins identified based on ≥ 70 % identity and ≥ 70 % coverage.

Predicted gene models were obtained from the JGI automated annotation pipeline, which integrates *ab initio* gene prediction with transcriptomic evidence. Expressed sequence tags (ESTs) derived from publicly available Illumina EST (MycoCosm Genome Interface) assemblies were aligned to the genomic regions using the RNNotator track with BLAT, confirming transcriptional support for predicted open reading frames (ORFs) (Martin et al. 2010). Key features such as transcript boundaries, protein domain architecture, and homologous sequence alignments were used to validate and interpret the predicted rhodopsin gene models. The transcript data are available in the JGI GOLD SRA explorer and NCBI’s SRA (Sequence Read Archive) database. The CDS region of our predicted modular rhodopsins were used as a query sequence in the NCBI’s tBLAST tool against concerned SRA data. Further, the hits obtained after tBLAST, were aligned with the CDS of the predicted modular rhodopsins using Clustal Omega (https://www.ebi.ac.uk/jdispatcher/msa/clustalo).

### Phylogenetic pattern of the new group of novel modular fungal rhodopsins

The modular rhodopsins retrieved from JGI, NCBI and UniProt database were aligned with different well-characterized rhodopsins, including BR, ChR2, and ASR. The newly retrieved sequences were aligned by the Clustal ω program of MEGA version 11 against the above-mentioned well-characterized protein sequences (Tamura et al. 2021). The output of the analysis was utilised for the phylogenetic analysis through MEGA version 11 software by the maximum likelihood approach based on the JTT matrix-based model using a thousand bootstrap replicates (Kumar et al. 2018).

### Alignment-based detection of essential residues in rhodopsin transmembrane (TM) helices and modular domains through sequence homology

The identified rhodopsins were subsequently examined for functionally conserved critical amino acid residues and retinal binding motifs. These motifs were used to identify and classify these novel modular rhodopsins. The well-characterized LR, BR (light-gated ion pump), ASR and ChR2 (light-gated ion channel) were chosen as the reference proteins. The alignment analysis was carried out using Clustal Omega Multiple Sequence Alignment tool (https://www.ebi.ac.uk/jdispatcher/msa/clustalo). The existence of the crucial conserved residues identified within the novel modular rhodopsins were visualized through the BioEdit software (Hall et al. 2011).

Further, to analyze the existence of a potential retinal chromophore synthesis pathway, the key enzymes associated with the carotenoid biosynthesis pathway in the *Blastocladiella emersonii* genome (Galindo-Solis and Fernandez 2022) were utilized as a query sequence in this study. The protein sequences from this collection were subjected to the BLASTp search against the NCBI and Mycosom databases. The key proteins/enzymes participating in the retinal synthesis pathway were analyzed using the STRING software 12.0 (https://string-db.org/) in *Aureobasidium pullulans*. The top 20 interacting partners were analysed using cytohubba via Cytoscape software 3.10.4 (https://cytoscape.org/).

The sequence alignment of the RPEL modular domains found in different fungi, was carried out using Clustal Omega Multiple Sequence Alignment tool (https://www.ebi.ac.uk/jdispatcher/msa/clustalo). Further, the interacting partners of the modular rhodopsin from *A. pullulans* (ApRh-RPEL) were analysed using the STRING software 12.0 (https://string-db.org/). The top 50 interacting partners were analysed using cytohubba via Cytoscape software 3.10.4 (https://cytoscape.org/).

### Structure-function determinants and homology modeling of the fungal rhodopsin domain: An *Ab initio* structure prediction, molecular docking and structural homology

*Ab initio* three-dimensional (3D) structure prediction was accomplished with the help of Alphafold2 Colab (David et al. 2022). Hydrogen atoms were added, and Gasteiger charges were computed using the dockprep program of UCSF Chimera for structure preparation prior to molecular docking. The 3D structure of all-*trans* retinal was retrieved from the PubChem database (CID: 638015) (Kim et al. 2025). The ligand structure was optimized by a structure minimization algorithm in UCSF Chimera. The coordinates of retinal binding pocket were as per the reference bacteriorhodopsin (PDB ID: 1KGB) close to the lysine residue of 7^th^ transmembrane helix for the formation of Schiff base bond. Possible interaction of all-*trans* retinal was predicted by molecular docking with help of UCSF Chimera and AutoDock Vina (Pettersen et al. 2004; Trott and Olson 2010). The resulting protein-ligand interactions were ranked based on the binding affinity scores. The top ranked poses were selected for interaction analysis via PyMOL (DeLano and Bromberg 2004).

Protein-ligand interaction profiler (PLIP) was used to study the possible interaction of retinal molecule and rhodopsin inside retinal binding pocket (Salentin et al. 2015). The Swiss-model was used for structural homology analysis. The structure was compared on the Swiss-model platform and comparative overlapping was achieved with help of UCSF Chimera (Bienert et al. 2017; Pettersen et al. 2004; Waterhouse et al. 2018). The secondary structure of the NAD(P) Rossmann fold modular domain was predicted using the PSIPRED tool (McGuffin et al. 2000).

The closest structural homologs were identified using the default algorithms of the SWISS-MODEL homology modeling platform. Structural validation was performed using ERRAT and PROCHECK available through the UCLA DOE Lab server (Colovos and Yeates 2008; Laskowski et al. 1996; Laskowski et al. 1993). ERRAT is known for its analysis of non-bonded atomic interactions. While PROCHECK analyse the quality of protein structure based on stereochemical quality of the given protein. The predicted structures were subsequently superimposed onto the closest well-characterized structural homologs using UCSF Chimera. Overlap of the retinal-binding pocket was confirmed through detailed structural visualization.

### Biocuration of protein-protein interactome (PPI) of newly identified multi-domain containing fungal rhodopsins

The protein-protein interactions of effector domain(s) in association with rhodopsins, i.e., RPEL, MCM, and NAD(P), were predicted by the STRING database (von Mering et al. 2003). The predicted protein-protein interactome obtained as an output, was subsequently used to generate the network via Cytoscape 3.7.2 (Saito et al. 2012; Shannon et al. 2003). The genome sequence of the ascomycete, *A. pullulans*, was downloaded from the NCBI database and was employed as a template for the identification of Biosynthetic Gene Cluster (BGCs) across its entire genome via antiSMASH (Blin et al. 2017) fungal version. The genome sequence was uploaded in FASTA format, and annotations were provided in GFF format. The default parameters were used for the detection of biosynthetic genes and other supporting proteins (enzymes, transporters and cluster borders). The regulatory and functional interactions of BGC genes were analyzed by protein-protein interaction (PPI) network in the STRING database with medium confidence score. The proteins identified in BGCs analysis was individually used as a query to predict interaction based on the literature studies, co-expression, neighbourhood, and text mining. The predicted PPI network was exported to Cytoscape v 3.10.3 (Saito et al. 2012) for further analysis. The genes were subjected to functional enrichment and were grouped accordingly.

To explore the light-based modulation of BGCs, the rhodopsin RPEL, a light-sensitive modular photoreceptor, was introduced in the interaction network. The objective of such inclusion was to monitor the possible metabolic flux modulation by light. The metabolic pathways related to terpenoid, carotenoid and sphingolipid biosynthesis were prioritised for the opto-synthetic modulation of biosynthetic compounds.

## Results

### Fungal genome mining reveals novel microbial modular rhodopsins featuring transmembrane helices and effector domains

The genome mining using the NCBI, and JGI Mycocosm databases led to the identification of four distinct types of novel modular rhodopsins from different fungal organisms. These fungal rhodopsins were characterized on the basis of the type of effector domain. The four distinct modular domains coupled with rhodopsin were: 1) RPEL motif; 2) MCM (Mini-chromosome maintenance); 3) NAD(P) Rossmann fold (Motif binding to NAD^+^ cofactor), and 4) GC-cAT (Guanylate cyclase coupled with carnitine o-acyltransferase) **(Fig. 1)**. Apart from the four distinct novel modular domains, a striking modular rhodopsin was identified in fungi, *Rhizoclosmastium globosum*, *Obelidium mucromatum* and *Gorgonomyces haynaldii*, comprising of two distinct GC (Guanylate cyclase)-coupled rhodopsins (Rh-GC) in single ORF (open reading frame) constitutively **(Fig. S1)**. Further, a unique fungal halorhodopsin from *Cladophialophora carrionii* was identified for the first time in this study **(Fig. S1)**.

**Fig. 1.**
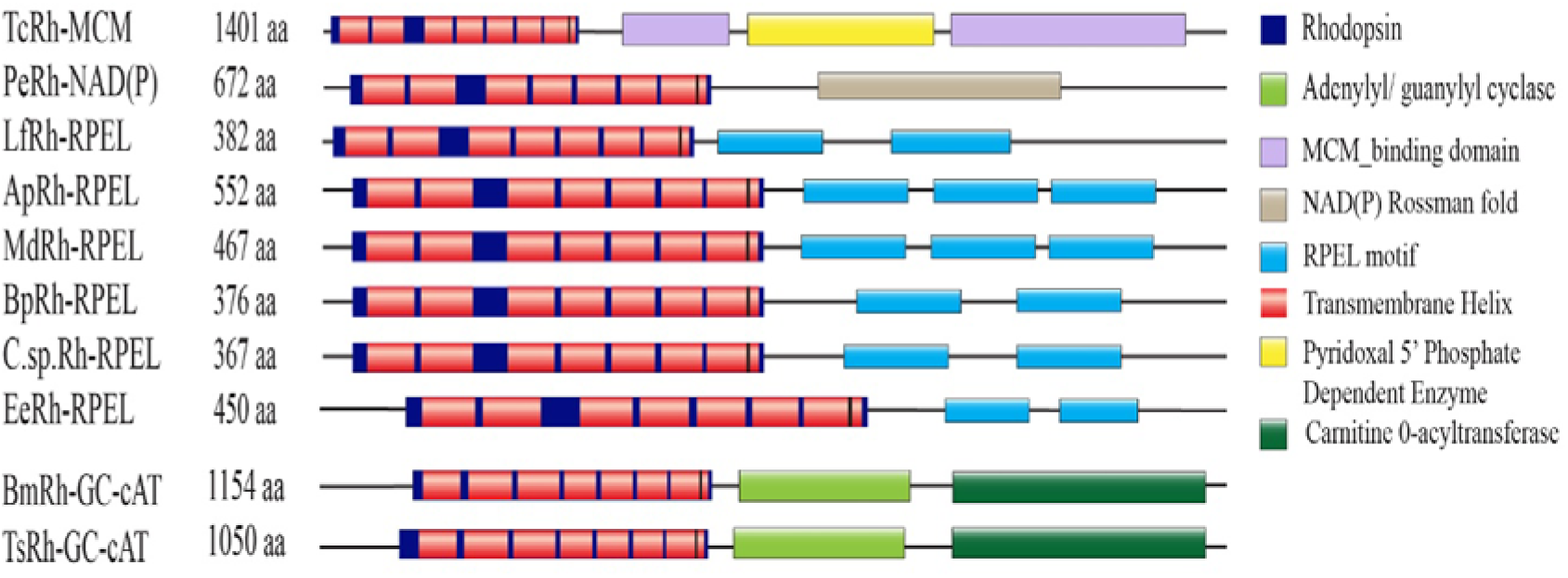
The schematic illustration depicts novel fungal rhodopsins with modular domains. Domain organisation of modular rhodopsin coupled with MCM (Minichromosome Maintenance) in *Testicularia cyperi* (TcRh-MCM), NAD(P) Rossmann Fold in *Pseudographis elatina* (PeRh-NAP(P)), RPEL repeats in *Lenthithecium fluviatile* (LfRh-RPEL), *Aureobasidium pullulans* (ApRh-RPEL), *Myriangium duriaei* (MdRh-RPEL), *Baudoinia panamericana* (BpRh-RPEL), *Capnodiales* sp. (C.sp.Rh-RPEL), guanylate cyclase (GC) coupled carnitine *O*-acyltransferase from (cAT) in *Boothiomyces macroporosus* (BmRh-GC-cAT), and GC coupled with cAT in *Terramyces subangulosus* (TsRh-GC-cAT). The blackline denotes the full-length protein, and domains are illustrated by geometric structures.

The current work focuses on the characterization of four novel modular rhodopsins using bioinformatics tools. The conserved domain analysis of these novel modular rhodopsins, conducted via the NCBI Conserved Domain Database (CD Search analysis), revealed the presence of the prominent seven-transmembrane helices (7TM), a distinguishing feature of rhodopsins. However, the twin modular rhodopsin (rhodopsin-GC) exhibited variability in the number of transmembrane helices, ranging from five to six, with additional flanking helices of unknown function **(Fig. S1)**. The accession number of the newly identified modular rhodopsins is listed in **Table 1**. The presence of the 7TM in the Rh-RPEL, Rh-MCM, Rh-NAD(P) Rossmann fold and Rh-GC-cAT, halorhodopsin was further confirmed using DeepTMHMM-1.0 bioinformatics tool. The in*-silico* tool successfully predicted the 7TM in each of the modular rhodopsins and halorhodopsins identified in this study. Based on the prediction, amino acids located within the transmembrane region, and extracellular and intracellular loops, the schematic representation of each of the 7TM of the modular rhodopsins were prepared **(Fig. S2)**. To verify that the predicted modular rhodopsins are expressed, we examined publicly available RNA-seq datasets and EST libraries from the NCBI SRA and JGI databases. *De novo* transcript assemblies using RNNotator and Illumina EST data revealed that each modular rhodopsin, Rh-RPEL, Rh-MCM, Rh-GC-cAT, and RhGC-RhGC was encoded by a single contiguous transcript. Regions that appeared fragmented in genome annotations were confirmed by matching EST-supported contiguous nucleotide stretches, that was further confirmed by JGI Genome Browser. The presence of complete or partial ESTs corresponding to the rhodopsin-effector fusion regions support the integrity and expression of these novel modular rhodopsins *in vivo* **(Fig. S3)**.

### Diversification of novel fungal modular rhodopsins uncovers unique evolved lineages

Brown (Brown 2004) identified fungal homologs of bacteriorhodopsins, classifying them into true rhodopsins (RDs), which preserve the retinal-binding lysine, and opsin-related proteins (ORPs), which lack this residue. In this study, we focused on a bacteriorhodopsin-like modular rhodopsin. To explore its evolutionary relationships, we concentrated on the RD subgroup and incorporated our sequence into Brown’s dataset for comparative phylogenetic analysis. The inclusion of ChR2 (channelrhodopsin), ASR (*Anabaena* sensory rhodopsin), and BR (bacteriorhodopsin) provides reference points for comparison with canonical microbial rhodopsins.

The Rh-RPEL (rhodopsin of the Rh-RPEL domain) clade is clearly separated from other rhodopsin types **(Fig. 2)**. Its tight clustering, supported by high bootstrap values, indicates functional conservation, while minor divergences suggest adaptation to specific fungal lineages or habitats. The ApRh-RPEL and MdRh-RPEL grouping, and the EeRh-RPEL and BpRh-RPEL grouping, indicate that they are very close relatives, likely orthologs or recent paralogs. Another strong sub-clade (supported by bootstrap values) includes true fungal rhodopsins (RDs) from *Ustilago*, *Cladonia*, *Zymoseptoria*, *Fusarium*, and *F. graminearum*. Their tight grouping reflects strong conservation among Ascomycota and Basidiomycota rhodopsins. *Pseudographis elatina* rhodopsin (PeRh) and *Botrytis* RD form another distinct sub-clade value, indicating shared ancestry among filamentous Ascomycetes and the evolution toward integrated photoregulatory roles. TcRh-MCM, ClacRh, *Zymoseptoria trictici* RD2, *Fusarium fujikuroi* RD, and *F. graminearum* RD2 form a strong sub-clade (mostly supported by 90-95 bootstrap values), suggesting they may share a more specific function. The BR and ASR branches out independently, suggesting they represent functionally or structurally divergent proteins. However, *C. carrionii* halorhodopsin (ClacRh) forms a distinct, deeply branching lineage within true fungal RDs. Its retention of the canonical NTQ chloride-pump motif hints at an evolutionary adaptation shaped by the organism’s environmental niche. To determine whether this represents a unique protein or a broader fungal group, a global BLASTp analysis was performed, revealing multiple fungal sequences with NTQ/DTQ motifs that cluster into a coherent, distinct clade **(Fig. S4)**. The *T. cyperi* rhodopsin (TcRh-MCM) and *Mycosarcoma maydis* (*Ustilago maydis*) rhodopsin (UstiRD2) form a well-supported sub-clade. The short branch length between them indicates high sequence conservation. TcRh-MCM carries an additional MCM-like domain, suggesting functional diversification toward light-regulated catalytic or signaling roles. The GC-cAT/ChR2 Clade (bottom section) is the most evolutionarily distinct group from the others, branching off earliest (lowest bootstrap support of 43 for the initial split with the rest of the tree). The Q8RUT8 ChR2 (channelrhodopsin from *C. reinhardtii*) is grouped with BmRh-GC-cAT and TsRh-GC-cAT. This suggests that these sequences, which are likely guanylyl-cyclase and channelrhodopsin-like enzymes, have diverged significantly from the large Rh-RPEL/RD group. The deep split indicates that this functional family separated from the others very early in the evolutionary history of these proteins. In conclusion, the tree highlights that the protein sequences fall into three or four major evolutionary families (clades), with the Rh-RPEL and RD sequences being the most numerous and closely related. On the other hand, the GC-cAT/ChR2 sequences represent the most ancient and divergent lineage among the proteins in this study. The accession numbers of the sequences used in the phylogenetic tree analysis are listed in **Table 2**.

**Fig. 2.**
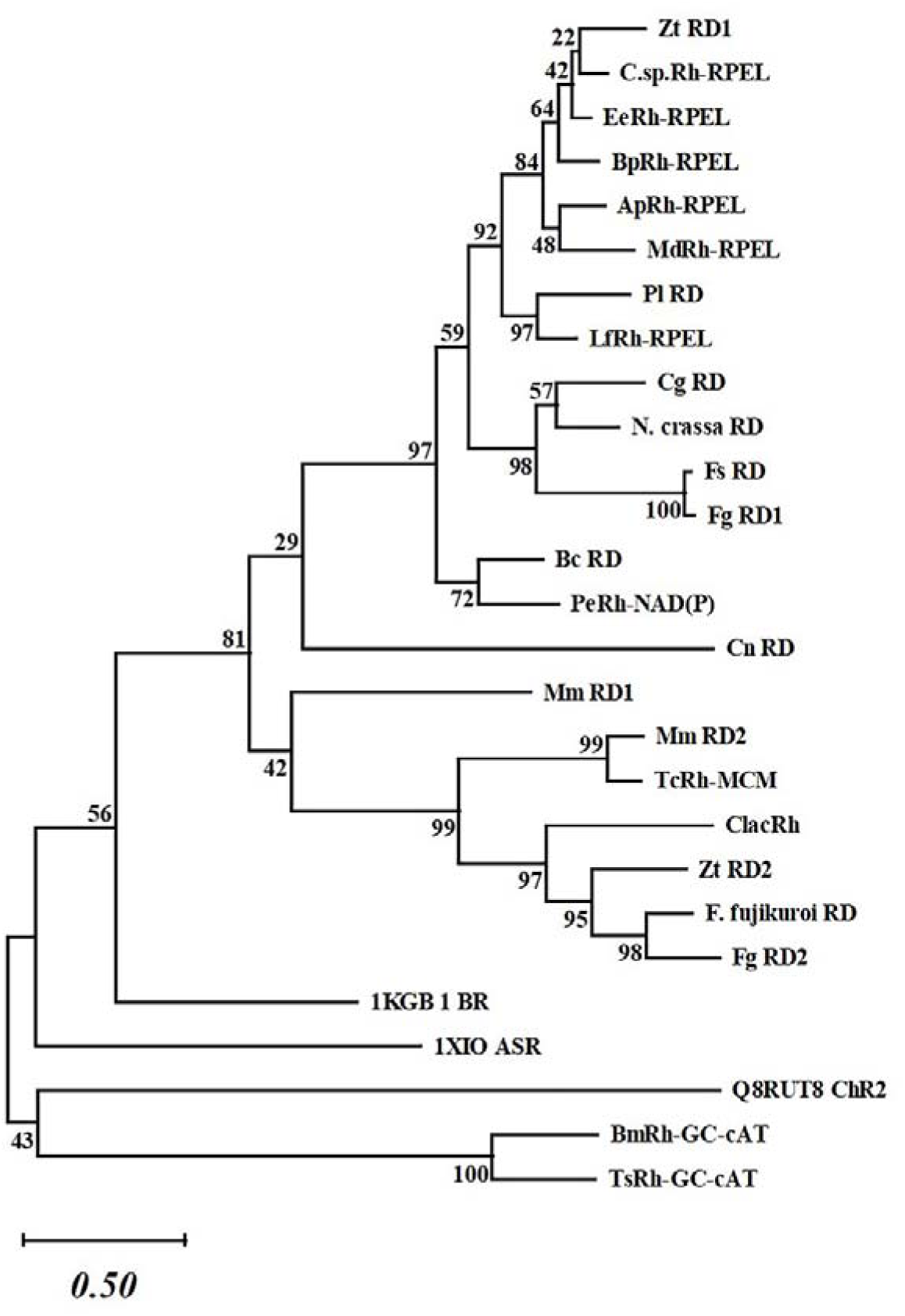
Phylogenetic analysis of novel fungal modular rhodopsins. The evolutionary diversification of microbial type rhodopsin was analysed using the maximum likelihood approach based on the JTT matrix-based model. It incorporates 1000 bootstrap values. The scale bar (0.5) at the bottom represents the branch length.

### Characterization of identified sequences for functional prediction of light-driven ion channels, sensory, and ion pumps based on conserved residues of the microbial rhodopsins

Rhodopsin activation and functions are mostly determined by the position of the absolute amino acid located close to the retinal binding site. Channelrhodopsin and bacteriorhodopsin are types of microbial rhodopsins that utilize retinal as a chromophore and experience light-induced structural changes; however, they vary in their proton pathways and functional results (Deisseroth and Hegemann 2017; Lanyi 2004; Nagel et al. 2003).

### Identification of novel fungal modular light-driven ion pump

The search for modular rhodopsins led to the identification of four distinct modular rhodopsins, each displaying unique domain architectures, as shown in **Fig. 1**. To investigate the structural conservation and functional relevance, the rhodopsin domains were aligned with well-characterized reference sequences, including BR, ASR, ChR2, and LR from the pathogenic fungus *Leptosphaeria maculans*. The phylogenetic analysis **(Fig. 2)** revealed that the novel modular domains, Rh-RPEL, Rh-MCM, and Rh-NAD(P), display greater evolutionary relatedness to BR and ASR compared to ChR2. In contrast, the Rh-GC-cAT shows more relatedness towards ChR2. Therefore, based on these results, sequence alignment studies were conducted, aligning the sequences of the modular domains Rh-RPEL, Rh-MCM, and Rh-NAD(P) with only BR and ASR. Subsequently, the Rh-GC-cAT sequences were aligned with ChR2.

The conserved residues essential for the photocycle have been highlighted in **Fig. 3A** and have been examined in the context of their major functional roles: (1) initiation of the photoisomerisation of retinal, which is covalently linked to Lys216 via a Schiff base; (2) facilitation of proton transfer from the Schiff base to Asp85; and (3) reprotonation of the Schiff base by Asp96 from the cytoplasmic side (Luecke et al. 2001). The amino acid residues, Arg 82, Glu194, and Glu204 are involved in facilitating the release of protons towards the extracellular side. The Asp212 residue aids in maintaining the correct electrostatic environment necessary for effective proton transfer. Depending on these amino acid residues, the rhodopsin domains were analyzed, and the details of the summary are in **Table 3** (Panzer et al. 2021). The boundary of the specific helices was taken according to the bacteriorhodopsin sequence, with a conserved lysine in the seventh helix. The identified rhodopsin sequences exhibit the same conserved triad residues in the third transmembrane helix as found in bacteriorhodopsin (BR), with the exception of TcRh, which contains a DTE motif instead of the canonical DTD seen in BR. At the position corresponding to G116 in BR, a site commonly used to distinguish fungal rhodopsins, we observed valine in PeRh and ApRh, leucine in TcRh, and isoleucine in other identified rhodopsin sequences. These residue variations may reflect functional divergence or evolutionary adaptations to specific ecological niches in fungal lineages **(Fig. 3A**; **Table 3)**. Overall, the strong conservation of key residues, including E194, E204, D212, and R82, suggests that these modular rhodopsins maintain a functional proton transfer mechanism. The presence of the conserved lysine residue in all sequences further supports their operational capability in light-induced activity.

**Fig. 3.**
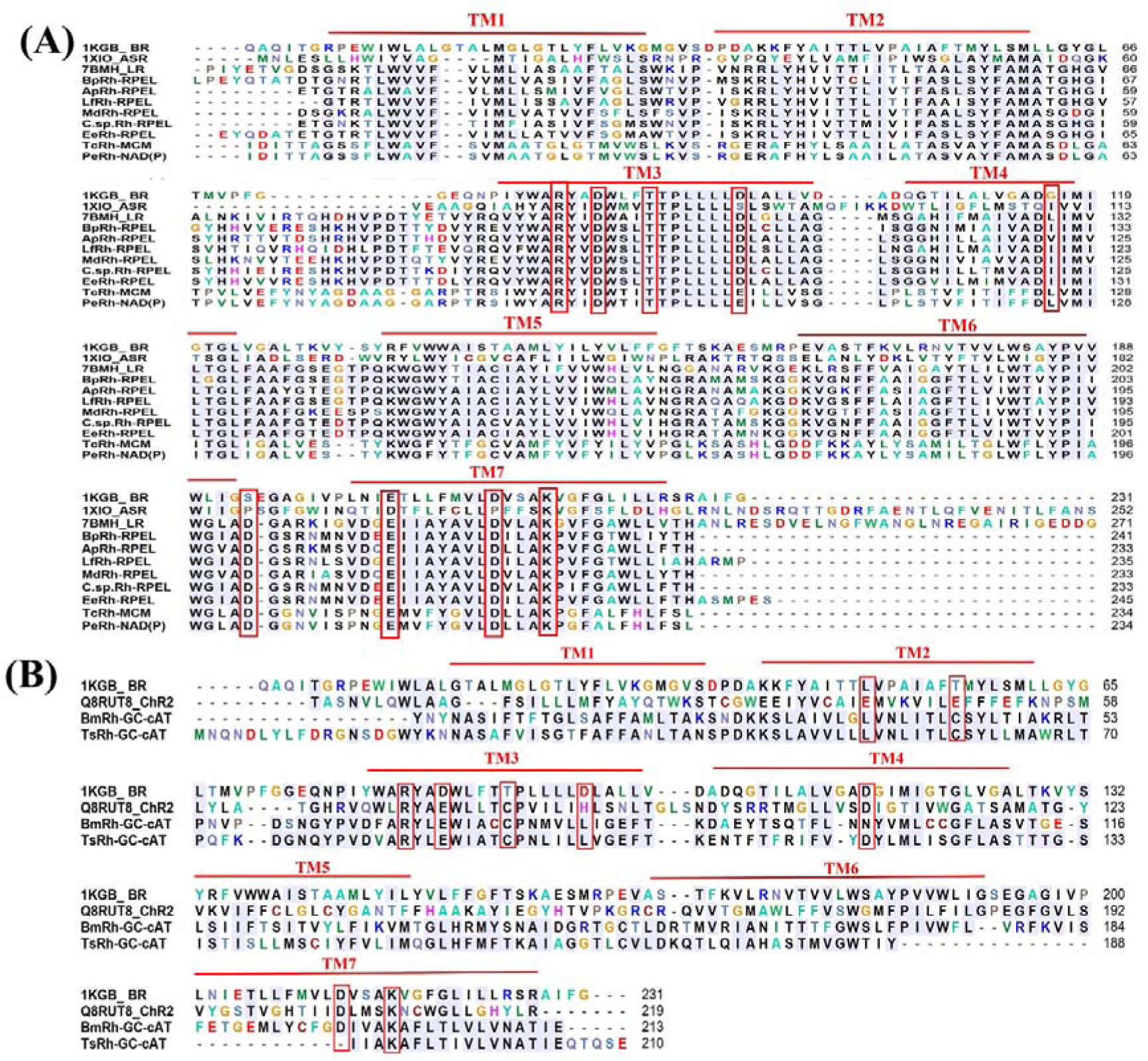
**(A)** Multiple sequence alignment of microbial rhodopsins from different fungi encoding the seven-transmembrane (7TM) domain and conserved retinal-binding lysine residue. Identical amino acids are marked in blue, and similar amino acids are highlighted in lighter green. The lysine residue is indicated by a red square. 1KGB: Bacteriorhodopsin, 1XIO: *Anabaena* sensory rhodopsin. **(B)** Comparative analysis of the putative fungal light-gated ion channels (BmRh and TsRh) and the well-characterized ChR2. The transmembrane helices (TM1–TM7) were illustrated as red bars. Critical residues are highlighted with red squares, which include the retinal-binding lysine, the proton donor and acceptor, the cysteine involved in forming the DC-gate through hydrogen bonding with the proton donor, and the arginine essential for primary proton translocation.

In addition to the modular rhodopsins, this study identified for the first time an HR (halorhodopsin)-like sequence, ClacRh, in *C. carrionii*, a pathogenic fungus known to cause human chromoblastomycosis. ClacRh exhibits overall similarity to BR but notably contains an NTQ motif, a signature feature of halorhodopsins, indicating potential differences in ion specificity and functional properties (Inoue et al. 2014).

### Identification of novel fungal modular channel-like sequences

A novel type of rhodopsin displays a distinctive arrangement of domains with the rhodopsin at the N-terminus, and guanylate cyclase coupled with carnitine *O*-acyltransferase at the C-terminus (Rh-GC-cAT). The Rh-GC-cAT modular rhodopsin was found to be encoded within a single open reading frame in the *B. macroporosus* (BmRh-GC-cAT) and *T. subangulosus* (TsRh-GC-cAT). The rhodopsin domains of BmRh and TsRh were analysed to evaluate their potential functionality. These domains were aligned with the well-characterized ChR2 and BR, which served as the reference rhodopsin. The key amino acid residues that are essential for the photocycle were examined, focusing on their primary roles: (1) the lysine that forms a covalent bond with retinal; (2) the counterion; (3) residues that maintain the proton acceptor environment; and (4) the DC-gate, located between helices three and four, which regulates channel function. Notably, the retinal-binding lysine in the seventh helix is conserved in both BmRh and TsRh, supporting their potential light-activated functionality **(Fig. 3B)**. BmRh possesses all seven conserved transmembrane helices typical of rhodopsins (**Fig. S2B)**, whereas TsRh lacks the seventh helix but retains the key lysine residue, suggesting a potentially altered retinal binding pocket, and it may be due to incomplete or poor-quality genome sequencing rather than true biological absence of the TM7 region. The aspartate residue at position 156 in ChR2 plays a key role in donating a proton to the retinal Schiff base (RSB) during re-protonation. This interaction is conserved in TsRh, but in BmRh it is replaced by asparagine (Asn) which may weaken cation coordination and affect the proton transfer efficiency **(Fig. 3B)**. Asp156 in ChR2 forms a hydrogen bond with Cys128, constituting the DC-gate that acts as a molecular switch for ion movement (Nack et al. 2010). Notably, Cys128 is conserved in the newly identified rhodopsin suggesting a potentially functional DC-gate mechanism **(Fig. 3B)**. Consequently, key functional residues in BmRh and TsRh, corresponding to those in ChR2, exhibit notable substitutions. These include E90N and E97Y substitutions that are crucial for pore gating. Additionally, BmRh displays another set of substitutions, D156N and H137L, that affect the internal proton donation and channel function **(Fig. 3B)**. Moreover, the presence of hydrophobic transmembrane domains enriched with aromatic residues may contribute to a positively charged microenvironment capable of supporting anion selectivity or alternative ligand interactions.

### Comparative structural-functional analysis of newly identified fungal rhodopsin

Structural superposition of LR with predicted light-driven ion-pump revealed a highly conserved seven-transmembrane helical architecture **(Fig. 4A)**. Ensemble superposition of representative proton-pumping rhodopsins aligned to LR revealed tight overlap of the seven-transmembrane (7-TM) helices, with an overall RMSD of 0.38 Å, indicating near-identical backbone organization across the group. Pairwise RMSD values, calculated using the PyMOL align function (cycles = 2, cutoff = 1.5 Å), ranged from 0.41 to 0.54 Å for most structures, indicating near-identical backbone organization. A modestly higher RMSD was observed for TcRh (0.85 Å), likely reflecting increased flexibility in loop or terminal regions rather than disruption of the core fold. In contrast, the structural alignment of LR with representative identified bacteriorhodopsin-like channelrhodopsin (BmRh and TsRh) resulted in a markedly higher overall RMSD (1.78 Å), reflecting substantial divergence consistent with functional specialization toward ion channel activity **(Fig. 4B)**. To further assess the conservation of the photochemical core, the three-dimensional structures of newly identified modular rhodopsins were compared with those of well-characterized rhodopsins from diverse systems. Consequently, the docking analyses revealed favorable binding energies for both all-trans retinal (−9.0 kcal/mol) and 11-cis retinal (−10.5 kcal/mol). Superposition of ApRh (brown) with ASR (cyan) revealed a conserved and overlapping retinal-binding pocket **(Fig. 4C)**. Homology modeling identified LmRh, a light-regulated proton pump, as the closest structural homolog of ApRh, consistent with reports of light-gated proton pump activity in *A. pullulans* (Panzer et al. 2021). Although the function of the extracellular loop (ECL) in fungal rhodopsins remains unclear, it is hypothesised to contribute to membrane stability. Structurally, the 7TM helix were superimposed over each other showing variance at the ECL except TcRh **(Fig. 4C)**. Further, analysis of the retinal binding pocket using PLIP confirmed the formation of a retinal Schiff base and revealed stabilizing hydrophobic interactions within the retinal binding pocket **(Fig. 4C)**. Rhodopsin (BmRh) of BmRh-GC-cAT were revealed to contain conserved residues making it closer to channelrhodopsin (ChR). Hence, we superimposed and monitored the position of retinal binding pocket of BmRh (red) and channelrhodopsin (green) by UCSF chimera. The ChR (cyan) and BmRh (grey) were superimposed to each other except the flanking region of the proteins. The retinal binding pocket was seen to be conserved **(Fig. 4D)**. In this study, the predicted structure scores ranged between 0.8 to 0.93, showing high confidence by AlphaFold. The structures were validated by ERRAT and PROCHECK, where the scores from ERRAT ranged from 97 % to 100 %. Analysis by PROCHECK demonstrated no errors, although warnings were found. However, all the structures passed the quality test and zero errors were found **(Table S1)**.

**Fig. 4.**
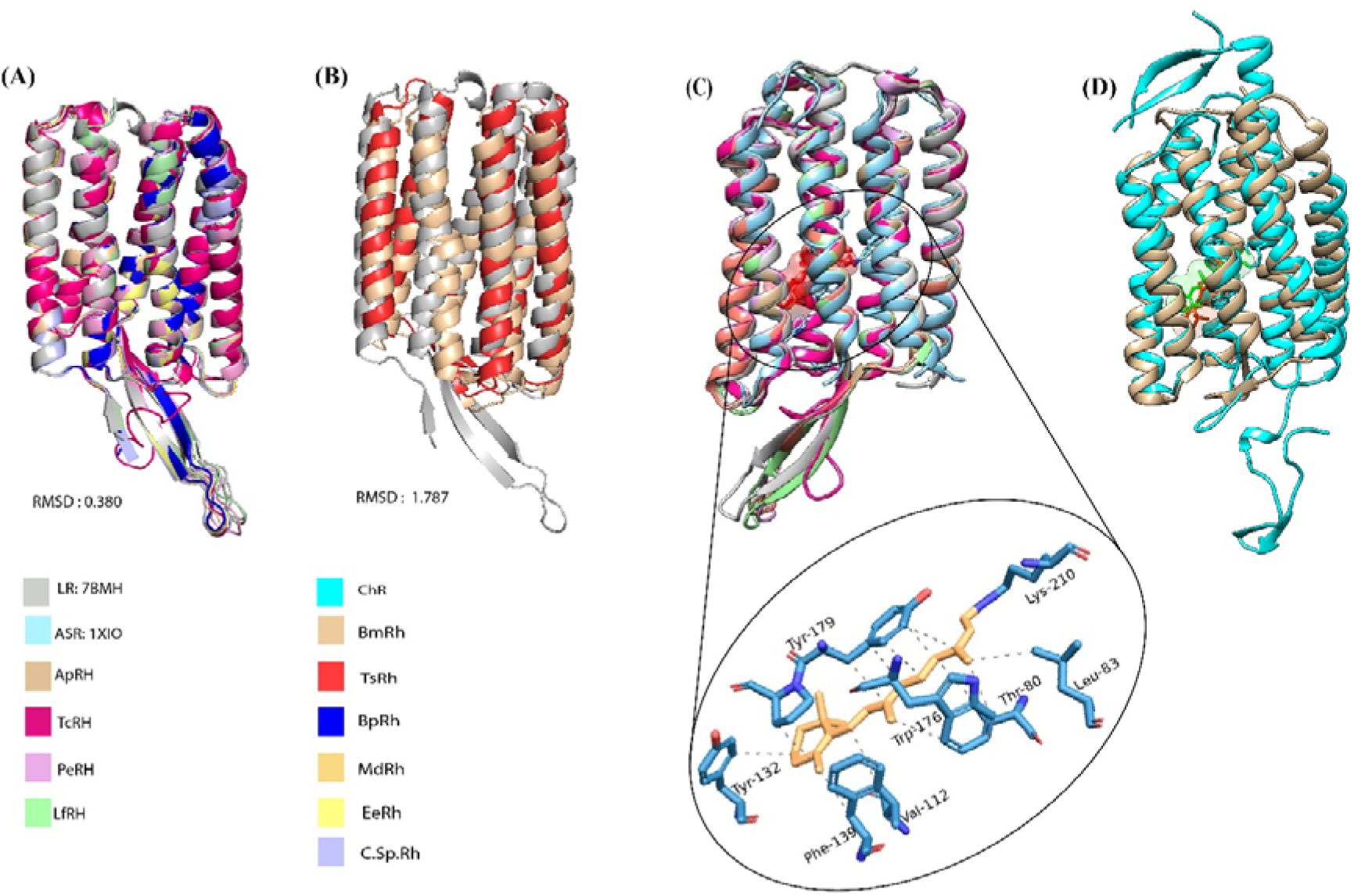
Structural analysis of modular rhodopsin with well-characterized bacteriorhodopsin, light-gated proton pump and channelrhodopsin (ion channel). **A)** ApRh (mustard colour) was docked with all-trans retinal (red) and super imposed with the sensory rhodopsin of *Anabaena* sensory rhodopsin (cyan). **B)** ApRh (mustard), along with other modular rhodopsins, were superimposed over *Anabaena* sensory rhodopsin ASR (PDB ID: 1XIO) and the light-gated proton pump of *Leptosphaeria maculans* (grey). **C)** All-trans retinal (orange) forming Schiff base bond with lysine residue (blue) of rhodopsin domain of the sensory rhodopsin. **D)** Superimposition of BmRh (grey) docked with all-trans retinal (red) and channelrhodopsin (cyan) show conserved retinal binding pocket.

### Modifications in the retinal-binding pockets of modular fungal rhodopsins resulted in a wider range of spectral tuning

Literature studies have shown that the analysis of the retinal binding pocket of microbial rhodopsins reveals differences in their absorption maxima. The variations in the absorption maxima are attributed to the amino acid composition within the retinal binding pocket (Hoffmann et al. 2006). Hence, in this study, the spectral tuning of the identified modular rhodopsin domains was targeted on the basis of amino acid residues within the retinal binding pocket with respect to BR, as shown in **Fig. 5**. Specifically, comparisons were made relative to the well-characterized BR and the variation at the 105^th^ position of proteorhodopsin (PR), which are summarized in **Fig. 5 and Table 4**. The key amino acid residues of the retinal binding pocket of BR and the identified novel modular domains were compared. The comparative study of the sequences of the rhodopsin with key spectral residues from BR predicted that the identified modular rhodopsins exhibit spectral shifts toward either blue or green light absorption, as summarized in **Table 4**. These findings are essential for the engineering of rhodopsins intended for use in optogenetics and photobiology. Our study found that the rhodopsin linked to the RPEL effector domain contains a conserved key spectral residue analogous to that in BR, corresponding to an absorption maximum (λmax) of approximately 568 nm. Thus, it can be engineered to produce red-shifted molecules, making them appropriate for optogenetic applications. Notably, substitutions such as D85E in BmRh and TsRh affect the stabilization of the protonated Schiff base, potentially causing shifts in the absorption maximum (λ_max_) depending on the surrounding molecular environment. Substitutions such as D85E in BmRh and TsRh affect the stabilization of the protonated Schiff base, potentially causing shifts in the absorption maximum (λ_max_) depending on the surrounding molecular environment. The D85N substitution in ClacRh reduces the counterion strength at the retinal Schiff base. Although generally considered a neutral change, it typically results in a red shift in the absorption spectrum. The T89C substitution in BmRh might introduce greater flexibility or a slightly altered hydrogen bonding pattern, resulting in fine-tuning. The T89 position in ClacRh introduces polarity similar to that in SRII, contributing to a λ_max_ shift of approximately 9-15 nm while preserving rhodopsin functionality. Leu93 of BR corresponding to position 105 in PR influences the size and steric properties of the retinal pocket, which determine whether the rhodopsin absorbs green (non-polar) and blue (polar) light. Similarly, our analysis revealed that smaller nonpolar side chains, Val in PeRh, Ile in ClacRh, and Met in BmRh, favour green light absorption, typically in the ∼520–540 nm range. It is suggested that ClacRh exhibits a naturally red-shifted wavelength, positioning it as the most effective optogenetic tool due to its superior light penetration and decreased phototoxicity **(Fig. 5 and Table 4)**.

**Fig. 5.**
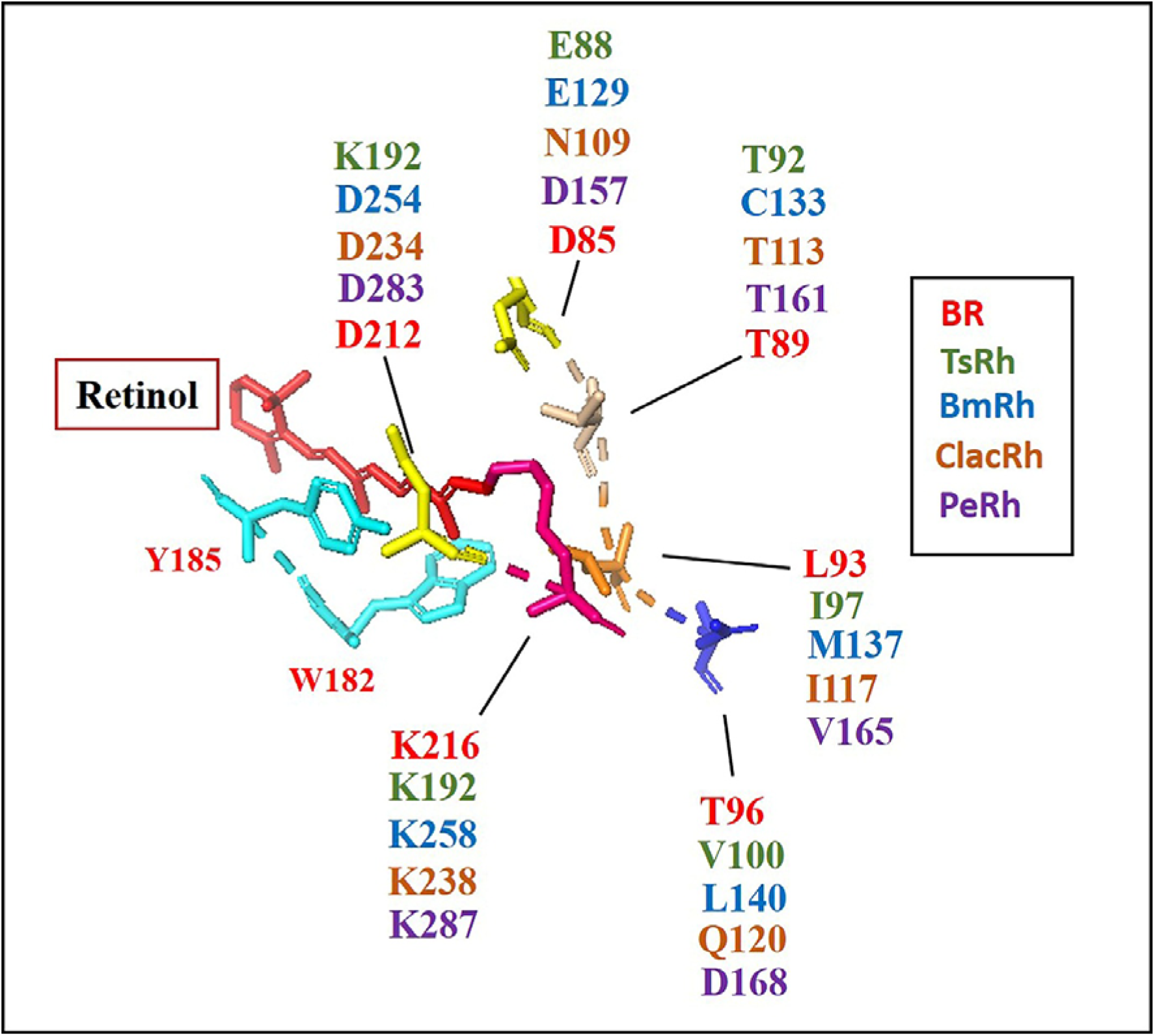
Comparative visualization of the functionally critical residues in the retinal-binding pocket of bacteriorhodopsin (BR) (PDB: 1KGB) and identified fungal rhodopsins. The analysis has been generated in PyMOL. The key surrounding residues of bacteriorhodopsin (BR) are labelled in red, which are compared with the residues found in TsRh (green), BmRh (blue), ClacRh (brown), and PeRh (purple).

### Presence of retinal biosynthesis pathway supports functional expression of modular rhodopsins in fungal system

In our analysis, we identified key genes involved in the β-carotene biosynthesis pathway required for retinal production in different fungal species. The retinal chromophore is essential for rhodopsin to function. It was found from the genome database of *A. pullulans* (Rh-RPEL)*, L. fluviatile* (Rh-RPEL)*, M. duriaei* (Rh-RPEL), and *T. cyperi* (Rh-MCM) that they contained all three enzyme-encoding genes involved in this pathway: bifunctional lycopene cyclase/phytoene synthase, phytoene desaturase, and carotenoid oxygenase. The key limiting enzymes found in other organisms that possess novel modular rhodopsins, such as those in other fungi, are summarized in **Table 5**. Therefore, the protein-protein interactome for the predicted carotenoid biosynthesis pathway of *A. pullulans*, that possesses a well-characterized genome, is depicted in **Fig. 6**. The curated network demonstrates significant connectivity among the enzymes involved in overlapping pathways (mevalonate pathway, carotenoid biosynthesis, and sterol metabolism). The interactome indicates that high-connectivity nodes, such as HMG-CoA reductase and GGPP synthase 1, might serve as regulatory bottlenecks. The presence of enzymes associated with carotenoid biosynthesis, including phytoene desaturase and lycopene beta-cyclase, suggests that *A. pullulans* possibly possesses a functional carotenoid biosynthesis pathway. The BLASTp analysis of the phytoene dehydrogenase sequence in *Capnodiales sp.* reveals similarities to the ATP-binding cassette transporter CGR1. The limited information available regarding the fungi *P. elatina*, *E. elasticus*, and *B. macroporosus* indicates that the study requires further experimental analysis.

**Fig. 6.**
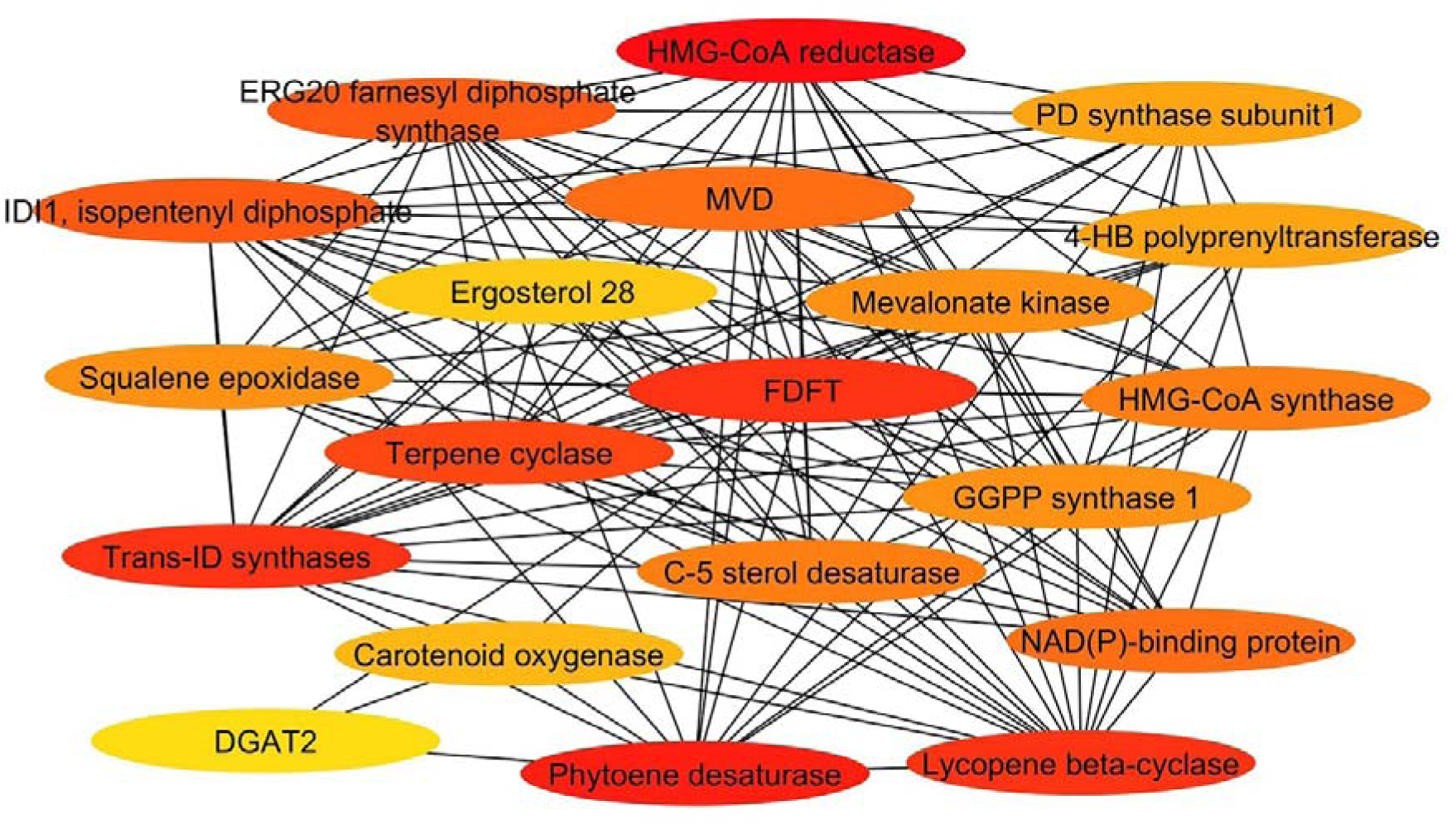
The protein-protein interaction (PPI) network of the carotenoid synthesis pathway in *Aureobasidium pullulans* predicted and visualized using STRING and Cytoscape, respectively. Nodes symbolize proteins, while edges indicate the interactions between them. The red nodes represent the proteins with the highest interactions within the PPI, where DGAT2: diacylglycerol O-acyltransferase 2, MVD: Mevalonate diphosphate decarboxylase, FDFT: Farnesyl-diphosphate farnesyltransferase, HMG CoA synthase: 3-Hydroxy-3-methylglutaryl-CoA synthase, GGPP synthase1: Geranylgeranyl diphosphate synthase 1, Trans ID synthase: Trans-isoprenyl diphosphate synthases, IDI1 isopentenyl diphosphate: dimethylallyl diphosphate isomerase, 4HB polyprenyltrasferase: 4-hydroxybenzoate polyprenyltransferase.

### The putative biological functions and optogenetic potential of the novel modular fungal rhodopsins

Amongst various effector domains associated with rhodopsin, the highlights were the RPEL repeats, NADP-binding Rossmann fold, MCM, and GC-cAT domains due to their putative functional relevance, which were evaluated for their significance. We performed functional assessments of these domains, where the protein-protein interaction (PPI) network (medium confidence score) analysis pinpointed their probable interacting partners and corresponding pathways.

The characterization of Rh-RPEL repeats is appealing because it may discover a wholly new class of type I modular rhodopsins. In the NCBI, BLASTp search, we found that its presence in the kingdom fungi is confined exclusively to the members of Ascomycota. It is demonstrated that G-Protein Coupled Receptors, GprM and GprJ in *Aspergillus fumigatus* are essential for regulating the cell wall integrity pathway, producing secondary metabolites, and influencing the organism’s virulence (Filho et al. 2020). Therefore, we hypothesize that a similar biological mechanism may exist that is perhaps regulated by the identified modular rhodopsin. The number of RPEL repeats influences the sensitivity of MRTF (Myocardin-related transcription factors) to changes in cellular actin dynamics (Watanabe et al. 2012). The proteins BpRh, MdRh, LfRh, C.sp.Rh and EeRh were predicted to include two RPEL repeat motifs, whereas ApRh features three RPEL repeat motifs according to the conserved domain analysis conducted through NCBI **(Fig. 1)**. The sequence homology analysis of RPEL repeats reveals a conserved RPxxxEL motif across most proteins. However, LfRh and MdRh show a substitution at the glutamate (E) position **(Fig. S5A)**. However the protein-protein interaction analysis for the ApRh-RPEL indicated its regulation in the G-actin polymerization biology. RPEL with proteins involved in protein folding and RNA modification, including key chaperones like Hsp70 and MAP kinases. These interactions influence gene expression and protein folding kinetics, which in turn affect the composition of secondary metabolites. Notably, RPEL’s association with proteins involved in sphingolipid metabolism suggests its role in modulating lipid-derived bioactive compound synthesis **(Fig. S5B & S6)**.

Rhodopsin-NADP-binding Rossmann fold-mediated optobiology indicates at a possible hybrid category of light-sensitive proteins that integrate light detection (through rhodopsin domains) with metabolic or redox functions through NAD(P)-binding Rossmann fold domains. This idea signifies a novel approach in synthetic biology and opto-biomanufacturing. Our study gives the first report to the scientific group that the rhodopsin-coupled NAD(P) Rossmann fold is naturally present in the non-lichenized fungus *P. elatina*.

However, the existing literature provides no information on the habitat of this fungus. The secondary structure analysis revealed a repeating β-α-β motif, characteristic of a classic Rossmann fold or a related Rossmannoid domain **(Fig. S7)**.

The fundamental biology of MCM proteins in fungi has been extensively investigated in several model organisms, yet there are considerable gaps in understanding regarding early-diverging fungi. For Chytridiomycota, Zygomycota, and Blastocladiomycota, not much is known about the regulation of MCM, which might exhibit distinct regulatory mechanisms. The complete sequence of Rh-MCM was examined using the NCBI conserved domain search program, revealing that it contains three consecutive domains: Rh-MCM, pyridoxal 5-phosphate (PLP)-dependent enzyme and MCM. For function prediction, we conducted protein-protein interaction (PPI) analysis using the full-length sequence as well as domain-specific analysis. However, we could not find specific PPIs related to MCM biology in the existing literature. To facilitate the potential optogenetic application of Rh-MCM, it is necessary to characterize the MCM biology of early-diverging fungus phyla. Currently, there is no experimental evidence directly demonstrating a physical or regulatory interaction between guanylate cyclase (GC) and carnitine acyltransferase (cAT) in fungi or other eukaryotic organisms. In *Candida albicans*, the expression of GC (CYC1) is enhanced during morphogenesis (Biswas et al. 2007) whereas cAT expression rises during changes in lipid metabolism, both of which may be triggered by nutrient scarcity (Rodaki et al. 2009). In this study, the following hypothesis is postulated: Guanylate cyclase activity influences cellular energy metabolism, thereby indirectly impacting the expression or activity of carnitine acyltransferase under conditions of nutrient stress. *B. macroporosus* is an uncommon and insufficiently described fungus making it challenging to investigate its putative rhodopsin-coupled guanylate cyclase with the cAT effector domain.

### Interaction of Biosynthetic Gene Cluster (BGC) with photoreceptors paves an unexplored way for opto-synthetic biotechnological avenues in the relevant fungal system

#### Light-mediated biosynthesis of fungal terpenoids and sphingolipids via rhodopsin-coupled RPEL module

The interaction between Biosynthetic Gene Cluster (BGC) proteins and photoreceptors in *A. pullulans* was investigated through a protein-protein interaction (PPI) network constructed using STRING software with medium confidence score. This analysis aimed to uncover potential light-regulated pathways influencing secondary metabolism. The predicted interactome analysis reveals that, within the terpenoid biosynthesis pathway, particularly enzymes like farnesyl pyrophosphate synthase (ERG20), are indirectly linked to the RPEL effector domain through interactions with protein folding complexes, transporter proteins, and tRNA synthases. This suggests a regulatory network where RPEL-mediated signaling may influence terpenoid production via intermediary cellular processes. These interacting partners act as crucial intermediaries linking the terpenoid biosynthesis pathway to the RPEL domain, which is coupled with rhodopsin. They may likely play significant roles in regulating protein synthesis, folding, and transport, suggesting a broader functional integration between light sensing and metabolic regulation. **(Fig. 7A)**. The RPEL motif is a G-actin binding element that regulates the activity of myocardin-related transcription factors (MRTF). These transcription factors possess several functions, interacting with other transcriptional regulators such as Hippo pathway effectors (YAP, TAZ) and TGFβ-controlled Smad proteins. Post actin polymerization, the MRTF along with its partner, serum response factor (SRF) directs gene expression for cytoskeletal remodelling, contractions, and several other processes (Miranda et al. 2021; Mouilleron et al. 2012). In fungi, actin is especially crucial for growth, morphogenesis and hyphal development (Hynes et al. 2011). Although the exact functioning and mechanistic basis of the RPEL-coupled rhodopsin’s regulation of terpenoid biosynthesis remain unclear, it is predicted that the light-controlled RPEL may direct changes in the fungal gene expression via upregulation/downregulation of transcription factors. The changes in gene expression alter the composition and content of fungal metabolites that may play influence the fungi’s pathogenicity, fungi-host interactions or motility (Galindo-Solis and Fernandez 2022).

**Fig. 7.**
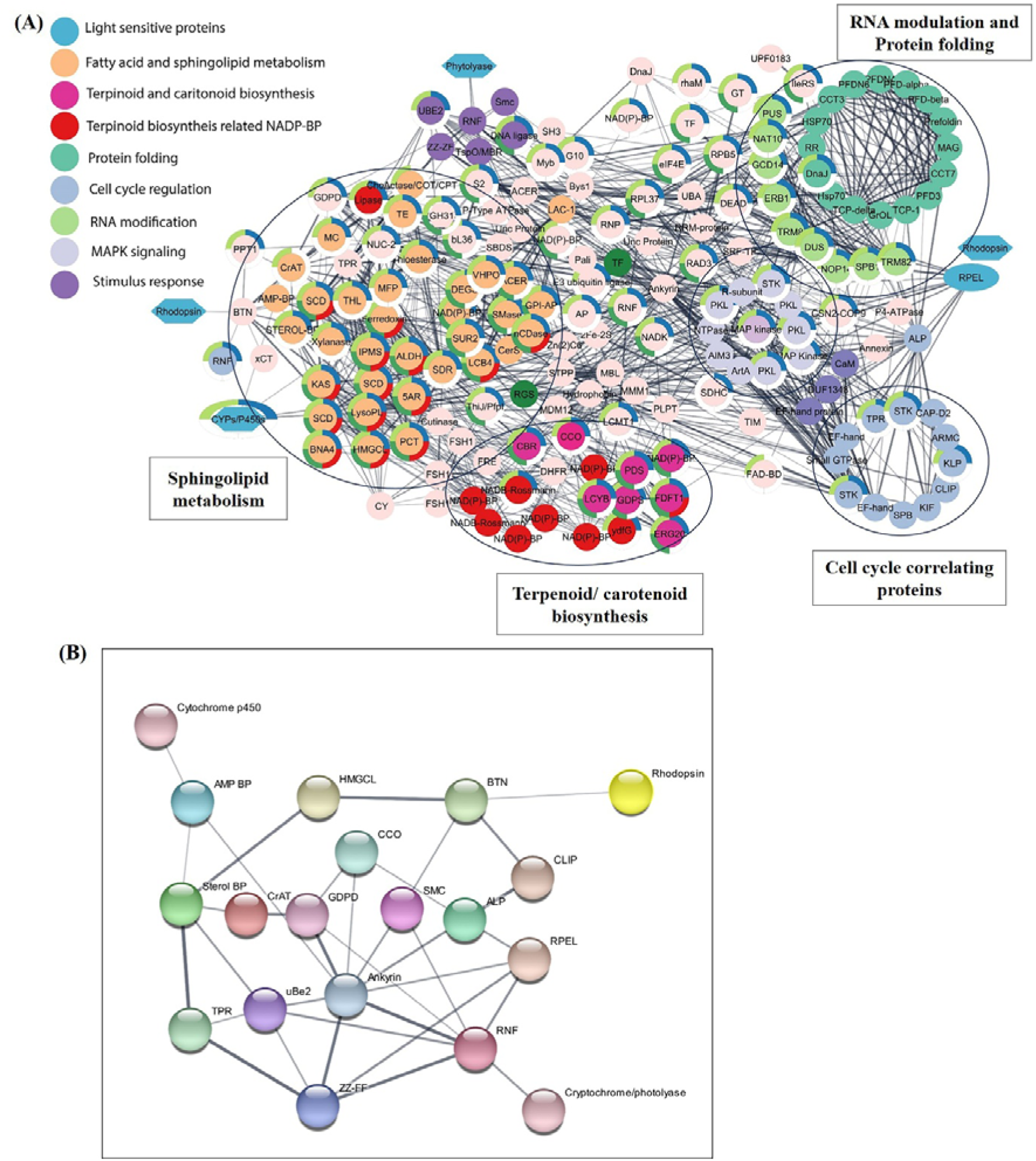
**(A)** Predicted opto-synthetic modulation of secondary metabolites in *Aureobasidium pullulans*. The interactome prepared via STRING software shows the interaction between light-sensitive proteins like rhodopsin, photolyase, cryptochrome, and the biosynthesis of secondary metabolites at different signaling pathways. **(B)** Manually curated protein-protein interactome prepared using STRING software, showing crosstalk between light-sensing proteins and their role in the regulation of the signaling cascade.

Similar to the terpenoid synthesis pathway, the PPI network predicts the interaction of the rhodopsin and RPEL with the actin-associated proteins (BTN, CLIP) regulating the function of sphingolipid biosynthetic pathway genes **(Fig. 7B)**. The constructed interactome predicts the potential crosstalk of the rhodopsin-regulated BTN, ALP and CLIP proteins with sphingolipid synthesis enzymes including carnitine acyltransferase (cAT) which is involved in the acyl-carnitine biosynthesis (Hynes et al. 2011). Further, other important photoreceptors such as LOV domain-containing proteins, photolyases and cryptochrome were also present in the developed network, which may directly modulate the biosynthesis of metabolites via light. The interactome highlights the possible interaction of these light-sensor molecules with the cytoskeletal proteins, ankyrin, ALP and even the RPEL domain. These cytoskeletal-associated proteins are anticipated to interact with several sphingolipid biosynthesis pathway proteins, including carnitine acyltransferase and sterol binding proteins.

## Discussion

Advances in *Chlamydomonas* photobiology enabled the first identification of modular rhodopsins; however, discoveries beyond algae remain limited. Recent metagenomic analyses reveal that fungi and flagellates also encode diverse rhodopsin-like genes, suggesting broader light-regulated pathways across microorganisms (Hou et al. 2012; Sineshchekov et al. 2002). Characterizing these modular rhodopsins is essential for understanding microbial responses to illumination and for uncovering light-dependent mechanisms that influence fungal bioactive production. In the present study, we targeted the mining of fungal genome databases using ChR2, ASR, BR, and HR as reference rhodopsins (Hou et al. 2012; Kawanabe and Kandori 2009; Verchere et al. 2017). We uncovered four previously unknown classes of modular fungal rhodopsins via bioinformatics tools. Furthermore, their modularity is defined by conserved effector motifs, including RPEL (RPxxxEL), MCM, NAD(P)-binding Rossmann folds, and GC-cAT, which reveal domain architectures.

These modular rhodopsins were identified across multiple fungal phyla, including Ascomycota, Basidiomycota, and Chytridiomycota, spanning both motile and non-motile members. Several non-motile Ascomycetes, including *B. panamericana*, *A. pullulans*, *L. fluviatile*, *E. elasticus*, *M. duriaei*, and *Capnodiales* spp., encode rhodopsins fused to an RPEL (RPxxxEL) motif. Because RPEL motifs mediate G-actin binding and regulate actin-dependent transcription factors such as MRTFs (Favot et al. 2005), their fusion to rhodopsins suggests a potential link between light sensing and cytoskeletal or transcriptional regulation. Two additional modular fusions were observed: a NAD(P)-binding Rossmann motif in *P. elatina* and MCM (minichromosome maintenance) motif in *T. cyperi*, which are non-motile fungal members of Ascomycota and Basidiomycota, respectively. The function of MCM-associated proteins (Corrochano 2019; Forsburg 2004; Kavakli et al. 2017; Sancar 2004; Yadav and Polasek-Sedlackova 2024) and NAD(P)-binding Rossmann fold is well established (Brzezinski 2020).

Given the central role of light in fungal development, including germination, conidiation, circadian entrainment, and reproductive transitions, these rhodopsin-coupled domains may represent light-responsive regulatory modules integrated into metabolic and genomic pathways. Their functional relevance, however, remains to be experimentally established. A fourth modular architecture was found in the zoosporic fungi *T. subangulosus* and *B. macroporosus*, which encode rhodopsins fused to a GC-cAT domain. Carnitine O-acyltransferase motifs are implicated in fungal metabolic and polyketide pathways, suggesting potential coupling between light sensing and metabolic regulation (Arishi et al. 2024; Keshavarz and Khalesi 2016). BmRh-GC-cAT carries substituted analogues of key proton-channeling residues known from ChR2, which may confer altered functional properties, although both the DC gate and the retinal-binding lysine remain conserved. This reinforces the notion that it likely maintains light sensitivity through retinal-bound lysine. While it might not function as a conventional ion channel, it appears to utilize proton-sensitive residues for conformational signaling. It advocates that BmRh and TsRh are likely derived from ChR-like ancestors to possibly fulfil photo-signaling functions. Due to its distinctive sequence, further experimental investigation is needed and could lead to engineering efforts aimed at improving its characteristics. As such, newly identified ion channels hold promise as optogenetic instruments for controlling novel biological pathways.

Light-driven signaling influences metabolic reprogramming and virulence in pathogenic fungi, highlighting its importance as a regulatory cue in fungal pathogenesis (Fuller et al. 2015). This is particularly relevant for *T. cyperi* and *C. carrionii*, pathogenic fungi identified in this study, which cause plant smut disease and human chromoblastomycosis, respectively (Badali et al. 2008; Kijpornyongpan et al. 2018). Photoreceptors including cryptochromes, phytochromes, white-collar proteins, and rhodopsins are also known to modulate growth and virulence in *Ustilago maydis* (Cerón-Bustamante et al. 2023), suggesting that rhodopsin-linked regulatory modules may contribute to pathogenic traits. The presence of Rh-MCM in *T. cyperi* and a chloride-pump like halorhodopsin in *C. carrionii* aligns with their placement in distinct phylogenetic clades and may reflect functional specialization associated with ecological adaptation. The adaptation of fungi to their habitats can affect the way they are distributed, specialized, and diversified, influencing their ecological functions and evolutionary paths. Its divergence suggests functional specialization: unlike proton-pumping RDs, ClacRh has residues consistent with light-driven chloride transport, confirming its identity as a photoactive pump. Fungal rhodopsins likely mirror the spectral plasticity seen in other microbial rhodopsins, where subtle changes in counterion residues or retinal-pocket amino acids, such as the colour-tuning positions in BR, SRII, or proteorhodopsin, can shift λ_max without altering core photochemical function (Kralj et al. 2008; Shimono et al. 2001).

Because rhodopsin activity depends on the retinal chromophore, which fungi generate from β-carotene via carotenoid oxygenases, we examined *A. pullulans* for the presence of carotenoid biosynthetic enzymes (Avalos et al. 2017; Yang et al. 2021). Our analysis showed that key carotenoid pathway components are encoded in *A. pullulans*, supporting the potential for endogenous retinal production and functional expression of the identified Rh-RPEL modular rhodopsin. The search for new modular rhodopsins is central to advancing opto-biotechnology, as light offers precise, tunable control over cellular processes and metabolite production (Bills and Gloer 2016; Chamkhi et al. 2021; Keller 2019; Kumar et al. 2023; Vaishnav and Demain 2011).

In this study, systems-level analysis revealed that fungal rhodopsins can interface with BGC-associated pathways, suggesting potential light-dependent regulation of metabolite synthesis. Our analysis suggests that in *A. pullulans* the Rh-RPEL fusion may connect to carotenoid, sphingolipid, and terpenoid metabolism, that are the key contributors to fungal secondary metabolite production. Consistent with this, *A. pullulans* encodes the full carotenoid-retinal biosynthetic machinery required for rhodopsin function and is known to alter metabolite output under different light conditions, such as enhanced pullulan production under blue light (Bozoudi and Tsaltas 2018; Iqbal et al. 2023; Prasongsuk et al. 2018). The predicted blue-light sensitivity of ApRh-RPEL aligns with these observations and suggests a role in photoregulated metabolic output. Beyond RPEL-containing sequences, the identification of Rh-MCM and Rh-NADP-binding Rossmann folding domains in pathogenic and saprophytic fungi suggests that light sensing may intersect with genome maintenance or redox-dependent pathways. The discovery of RhGC-RhGC tandem fusions in *R. globosum* and a novel halorhodopsin in *C. carrionii* further expands the functional space of fungal rhodopsins and raises the possibility of specialized roles in environmental adaptation or virulence. The identification of channelrhodopsin-like sequences in chytrid fungi (*B. macroporosus* and *T. subangulosus*) underscores the evolutionary breadth of light-responsive systems in the fungal kingdom. Together, these findings reveal a previously unappreciated diversity of modular fungal rhodopsins and suggest that they may serve as opto-synthetic regulators of metabolic, developmental, and potentially pathogenic processes. Future biochemical and genetic studies will be essential to determine their photochemical properties, domain-specific functions, and applicability as tools for biological, optogenetics, or light-controlled biomanufacturing.

## Conclusions

This study uncovers an unexpectedly diverse repertoire of modular rhodopsins in fungi, revealing light-sensing cores fused to regulatory domains previously unknown in this kingdom and other organisms. These modular rhodopsins retain marks of functional microbial rhodopsins including conserved retinal-binding motifs, the canonical Schiff-base lysine in TM7, and putative seven-transmembrane architectures supporting their potential as light-responsive signaling modules. Although several sequences share limited similarity with known ion channels or pumps. It suggests that they represent evolutionarily divergent rhodopsin families with distinct functional repertoires. Predictive spectral analyses indicate that many newly identified rhodopsins could absorb predominantly in the green region, while specific substitutions in *C. carrionii* halorhodopsin and *P. elatina* rhodopsin suggest additional spectral tuning adaptations. Systems biology analysis along with curated BGC associations further reveals that the ApRh-RPEL fusion in *A. pullulans* may interface with pathways governing terpenoid, sphingolipid, and carotenoid biosynthesis, pointing toward a possible link between light perception and secondary-metabolite regulation. The integrity of all predicted rhodopsin-effector fusions is supported by EST and transcriptomic evidence, strengthening the confidence in their biological relevance. Collectively, these findings expand the known diversity of fungal rhodopsins and establish a conceptual framework for understanding their potential biological, optogenetic, opto-synthetic, and biotechnological utility. Future experimental validation will be essential to elucidate their mechanistic roles and harness their light-dependent regulatory capacities to advance optogenetic toolkits and fungal bioengineering applications.

## Glossary

**Fungal modular rhodopsin:** A Type-I microbial rhodopsin fused to regulatory or catalytic effector domains, enabling light-dependent modulation of downstream cellular processes.

**Twin rhodopsins**: It is a tandem rhodopsin protein in which two rhodopsin domains, each typically containing seven transmembrane helices and a retinal-binding lysine

**Biosynthetic Gene Cluster (BGC):** A biosynthetic gene cluster is a short stretch of DNA where several neighboring genes work together to make one specific natural product (such as an antibiotic, pigment, or toxin). All the key enzymes, regulators, and often transporters for that molecule are encoded side-by-side in this cluster, so they can be turned on and off as a unit.

**Opto-synthetic biology:** An interdisciplinary field that integrates optogenetics (light-controlled proteins, such as rhodopsins, LOV, CRY2/CIB, PhyB/PIF, etc.) with synthetic biology to design light-controlled metabolic pathways that can be turned on, off, or tuned simply by changing the light wavelength, intensity, or timing.

**Rossmann fold:** It is a common protein structural motif used for binding nucleotide cofactors such as NAD□, NADP, and FAD.

**RPEL motif**: It is also known as the RPXXXEL motif, a short, conserved peptide sequence that mediates high-affinity binding to G-actin (globular actin).

## Acknowledgements

The research grants (ECR/2017/000354 & CRG/2021/003158) from ANRF/SERB is duly acknowledged. Alka Kumari and Abhishek Kumar are the recipients of the Junior Research Fellowship from UGC (JRF) and ICMR (JRF), respectively. We are thankful to all the lab members for their kind support.

## Author contributions

**Suneel Kateriya:** Conceptualisation, Data curation and Methodology, Supervision, Validation, Funding acquisition, Writing-review and editing. **Alka Kumari:** Formal analysis, Investigation, Writing-original draft, Methodology. **Abhishek Kumar:** Formal analysis, Methodology. **Komal Sharma:** Methodology. **Swaroop Ranjan Pati:** Methodology. **Shilpa Mohanty:** Methodology, Writing-review and editing.

## Funding

We are thankful to ANRF/SERB, Government of India for granting ECR (ECR/2017/000354) and CRG (CRG/2021/003158) research projects.

## Data availability statement

Data is available on request to the corresponding author.

## Declarations

### Competing interests

The authors declare that they have no known competing financial interests or personal relationships that could have appeared to influence the work reported in this paper.

## Notes

### Competing Interest Statement

The authors have declared no competing interest.

## References

1. Altschul SF, Gish W, Miller W, Myers EW, Lipman DJ (1990) Basic local alignment search tool. J Mol Biol 215:403–410. 10.1016/S0022-2836(05)80360-2

2. Arishi AA, Shang Z, Lacey E, Crombie A, Vuong D, Li H, Bracegirdle J, Turner P, Lewis W, Flematti GR, Piggott AM, Chooi YH (2024) Discovery and heterologous biosynthesis of glycosylated polyketide luteodienoside A reveals unprecedented glucinol-mediated product offloading by a fungal carnitine O-acyltransferase domain. Chem Sci 15:3349–3356. 10.1039/d3sc05008d

3. Avalos J, Pardo-Medina J, Parra-Rivero O, Ruger-Herreros M, Rodriguez-Ortiz R, Hornero-Mendez D, Limon MC (2017) Carotenoid Biosynthesis in *Fusarium*. J Fungi (Basel) 3 10.3390/jof3030039

4. Avelar GM, Schumacher RI, Zaini PA, Leonard G, Richards TA, Gomes SL (2014) A rhodopsin-guanylyl cyclase gene fusion functions in visual perception in a fungus. Curr Biol 24:1234–1240. 10.1016/j.cub.2014.04.009

5. Badali H, Gueidan C, Najafzadeh MJ, Bonifaz A, van den Ende AH, de Hoog GS (2008) Biodiversity of the genus *Cladophialophora*. Stud Mycol 61:175–191. 10.3114/sim.2008.61.18

6. Bienert S, Waterhouse A, de Beer TA, Tauriello G, Studer G, Bordoli L, Schwede T (2017) The SWISS-MODEL Repository-new features and functionality. Nucleic Acids Res 45:D313–D319. 10.1093/nar/gkw1132

7. Bills GF, Gloer JB (2016) Biologically Active Secondary Metabolites from the Fungi. Microbiol Spectr 4 10.1128/microbiolspec.FUNK-0009-2016

8. Biswas S, Van Dijck P, Datta A (2007) Environmental sensing and signal transduction pathways regulating morphopathogenic determinants of *Candida albicans*. Microbiol Mol Biol Rev 71:348–376. 10.1128/MMBR.00009-06

9. Blin K, Medema MH, Kottmann R, Lee SY, Weber T (2017) The antiSMASH database, a comprehensive database of microbial secondary metabolite biosynthetic gene clusters. Nucleic Acids Res 45:D555–D559. 10.1093/nar/gkw960

10. Bozoudi D, Tsaltas D (2018) The Multiple and Versatile Roles of *Aureobasidium pullulans* in the Vitivinicultural Sector. Fermentation 4. 10.3390/fermentation4040085

11. Brown LS (2004) Fungal rhodopsins and opsin-related proteins: eukaryotic homologues of bacteriorhodopsin with unknown functions. Photochem Photobiol Sci 3:555–565. 10.1039/b315527g

12. Brzezinski K (2020) S-adenosyl-l-homocysteine Hydrolase: A Structural Perspective on the Enzyme with Two Rossmann-Fold Domains. Biomolecules 10. 10.3390/biom10121682

13. Cerón-Bustamante M, Balducci E, Beccari G, Nicholson P, Covarelli L, Benincasa P (2023) Effect of light spectra on cereal fungal pathogens, a review. Fungal Biology Reviews 43. 10.1016/j.fbr.2022.10.004

14. Chamkhi I, Benali T, Aanniz T, El Menyiy N, Guaouguaou FE, El Omari N, El-Shazly M, Zengin G, Bouyahya A (2021) Plant-microbial interaction: The mechanism and the application of microbial elicitor induced secondary metabolites biosynthesis in medicinal plants. Plant Physiol Biochem 167:269–295. 10.1016/j.plaphy.2021.08.001

15. Colovos C, Yeates TO (2008) Verification of protein structures: Patterns of nonbonded atomic interactions. Protein Science 2:1511–1519. 10.1002/pro.5560020916

16. Corrochano LM (2019) Light in the Fungal World: From Photoreception to Gene Transcription and Beyond. Annu Rev Genet 53:149–170. 10.1146/annurev-genet-120417-031415

17. David A, Islam S, Tankhilevich E, Sternberg MJE (2022) The AlphaFold Database of Protein Structures: A Biologist’s Guide. J Mol Biol 434:167336. 10.1016/j.jmb.2021.167336

18. Deisseroth K, Hegemann P (2017) The form and function of channelrhodopsin. Science 357. 10.1126/science.aan5544

19. DeLano WL, Bromberg SJDSL (2004) PyMOL user’s guide. 629

20. Emiliani V, Entcheva E, Hedrich R, Hegemann P, Konrad KR, Luscher C, Mahn M, Pan ZH, Sims RR, Vierock J, Yizhar O (2022) Optogenetics for light control of biological systems. Nat Rev Methods Primers 2. 10.1038/s43586-022-00136-4

21. Ernst OP, Lodowski DT, Elstner M, Hegemann P, Brown LS, Kandori H (2014) Microbial and animal rhodopsins: structures, functions, and molecular mechanisms. Chem Rev 114:126–163. 10.1021/cr4003769

22. Favot L, Gillingwater M, Scott C, Kemp PR (2005) Overexpression of a family of RPEL proteins modifies cell shape. FEBS Lett 579:100–104. 10.1016/j.febslet.2004.11.054

23. Filho A, Brancini GTP, de Castro PA, Valero C, Ferreira Filho JA, Silva LP, Rocha MC, Malavazi I, Pontes JGM, Fill T, Silva RN, Almeida F, Steenwyk JL, Rokas A, Dos Reis TF, Ries LNA, Goldman GH (2020) Aspergillus fumigatus G-Protein Coupled Receptors GprM and GprJ Are Important for the Regulation of the Cell Wall Integrity Pathway, Secondary Metabolite Production, and Virulence. mBio 11. 10.1128/mBio.02458-20

24. Forsburg SL (2004) Eukaryotic MCM proteins: beyond replication initiation. Microbiol Mol Biol Rev 68:109–131. 10.1128/MMBR.68.1.109-131.2004

25. Fuller KK, Loros JJ, Dunlap JC (2015) Fungal photobiology: visible light as a signal for stress, space and time. Curr Genet 61:275–288. 10.1007/s00294-014-0451-0

26. Furutani Y, Bezerra AG, Jr., Waschuk S, Sumii M, Brown LS, Kandori H (2004) FTIR spectroscopy of the K photointermediate of Neurospora rhodopsin: structural changes of the retinal, protein, and water molecules after photoisomerization. Biochemistry 43:9636–9646. 10.1021/bi049158c

27. Galindo-Solis JM, Fernandez FJ (2022) Endophytic Fungal Terpenoids: Natural Role and Bioactivities. Microorganisms 10. 10.3390/microorganisms10020339

28. Hall T, Biosciences I, Carlsbad CJGbb (2011) BioEdit: an important software for molecular biology. 2:60–61.

29. Hallgren J, Tsirigos KD, Pedersen MD, Almagro Armenteros JJ, Marcatili P, Nielsen H, Krogh A, Winther O (2022) DeepTMHMM predicts alpha and beta transmembrane proteins using deep neural networks bioRxiv. 10.1101/2022.04.08.487609

30. Hoffmann M, Wanko M, Strodel P, Konig PH, Frauenheim T, Schulten K, Thiel W, Tajkhorshid E, Elstner M (2006) Color tuning in rhodopsins: the mechanism for the spectral shift between bacteriorhodopsin and sensory rhodopsin II. J Am Chem Soc 128:10808–10818. 10.1021/ja062082i

31. Hou SY, Govorunova EG, Ntefidou M, Lane CE, Spudich EN, Sineshchekov OA, Spudich JL (2012) Diversity of *Chlamydomonas* channelrhodopsins. Photochem Photobiol 88:119–128. 10.1111/j.1751-1097.2011.01027.x

32. Hynes MJ, Murray SL, Andrianopoulos A, Davis MA (2011) Role of carnitine acetyltransferases in acetyl coenzyme A metabolism in *Aspergillus nidulans*. Eukaryot Cell 10:547–555. 10.1128/EC.00295-10

33. Inoue K, Koua FH, Kato Y, Abe-Yoshizumi R, Kandori H (2014) Spectroscopic study of a light-driven chloride ion pump from marine bacteria. J Phys Chem B 118:11190–11199. 10.1021/jp507219q

34. Inoue K, Ono H, Abe-Yoshizumi R, Yoshizawa S, Ito H, Kogure K, Kandori H (2013) A light-driven sodium ion pump in marine bacteria. Nat Commun 4:1678. 10.1038/ncomms2689

35. Iqbal M, Broberg A, Andreasson E, Stenberg JA (2023) Biocontrol Potential of Beneficial Fungus *Aureobasidium pullulans* Against *Botrytis cinerea* and *Colletotrichum acutatum*. Phytopathology 113:1428–1438. 10.1094/PHYTO-02-23-0067-R

36. Kandori H (2020) Biophysics of rhodopsins and optogenetics. Biophys Rev 12:355–361. 10.1007/s12551-020-00645-0

37. Kandori H, Inoue K, Tsunoda SP (2018) Light-Driven Sodium-Pumping Rhodopsin: A New Concept of Active Transport. Chem Rev 118:10646–10658. 10.1021/acs.chemrev.7b00548

38. Kato HE, Inoue K, Abe-Yoshizumi R, Kato Y, Ono H, Konno M, Hososhima S, Ishizuka T, Hoque MR, Kunitomo H, Ito J, Yoshizawa S, Yamashita K, Takemoto M, Nishizawa T, Taniguchi R, Kogure K, Maturana AD, Iino Y, Yawo H, Ishitani R, Kandori H, Nureki O (2015) Structural basis for Na(+) transport mechanism by a light-driven Na(+) pump. Nature 521:48–53. 10.1038/nature14322

39. Kavakli IH, Baris I, Tardu M, Gul S, Oner H, Cal S, Bulut S, Yarparvar D, Berkel C, Ustaoglu P, Aydin C (2017) The Photolyase/Cryptochrome Family of Proteins as DNA Repair Enzymes and Transcriptional Repressors. Photochem Photobiol 93:93–103. 10.1111/php.12669

40. Kawanabe A, Kandori H (2009) Photoreactions and structural changes of anabaena sensory rhodopsin. Sensors (Basel) 9:9741–9804. 10.3390/s91209741

41. Keller NP (2019) Fungal secondary metabolism: regulation, function and drug discovery. Nat Rev Microbiol 17:167–180. 10.1038/s41579-018-0121-1

42. Keshavarz B, Khalesi M (2016) Trichoderma reesei, a superior cellulase source for industrial applications. Biofuels 7:713–721. 10.1080/17597269.2016.1192444

43. Kijpornyongpan T, Mondo SJ, Barry K, Sandor L, Lee J, Lipzen A, Pangilinan J, LaButti K, Hainaut M, Henrissat B, Grigoriev IV, Spatafora JW, Aime MC (2018) Broad Genomic Sampling Reveals a Smut Pathogenic Ancestry of the Fungal Clade Ustilaginomycotina. Mol Biol Evol 35:1840–1854. 10.1093/molbev/msy072

44. Kim S, Chen J, Cheng T, Gindulyte A, He J, He S, Li Q, Shoemaker BA, Thiessen PA, Yu B, Zaslavsky L, Zhang J, Bolton EE (2025) PubChem 2025 update. Nucleic Acids Res 53:D1516–D1525. 10.1093/nar/gkae1059

45. Konno M, Kato Y, Kato HE, Inoue K, Nureki O, Kandori H (2016) Mutant of a Light-Driven Sodium Ion Pump Can Transport Cesium Ions. J Phys Chem Lett 7:51–55. 10.1021/acs.jpclett.5b02385

46. Kralj JM, Bergo VB, Amsden JJ, Spudich EN, Spudich JL, Rothschild KJ (2008) Protonation state of Glu142 differs in the green- and blue-absorbing variants of proteorhodopsin. Biochemistry 47:3447–3453. 10.1021/bi7018964

47. Kulbay M, Tuli N, Akdag A, Kahn Ali S, Qian CX (2024) Optogenetics and Targeted Gene Therapy for Retinal Diseases: Unravelling the Fundamentals, Applications, and Future Perspectives. J Clin Med 1310.3390/jcm13144224

48. Kumar A, Baldia A, Rajput D, Kateriya S, Babu V, Dubey KK (2023) Multiomics and optobiotechnological approaches for the development of microalgal strain for production of aviation biofuel and biorefinery. Bioresour Technol 369:128457. 10.1016/j.biortech.2022.128457

49. Kumar S, Stecher G, Li M, Knyaz C, Tamura K (2018) MEGA X: Molecular Evolutionary Genetics Analysis across Computing Platforms. Mol Biol Evol 35:1547–1549. 10.1093/molbev/msy096

50. Lanyi JK (2004) Bacteriorhodopsin. Annu Rev Physiol 66:665–688. 10.1146/annurev.physiol.66.032102.150049

51. Laskowski R, Rullmann JA, MacArthur M, Kaptein R, Thornton J (1996) AQUA and PROCHECK-NMR: Programs for checking the quality of protein structures solved by NMR. Journal of Biomolecular NMR 810.1007/bf00228148

52. Laskowski RA, MacArthur MW, Moss DS, Thornton JM (1993) PROCHECK: a program to check the stereochemical quality of protein structures. J Appl Crystallogr 26:283–291. 10.1107/s0021889892009944

53. Liu RS, Colmenares LU (2003) The molecular basis for the high photosensitivity of rhodopsin. Proc Natl Acad Sci U S A 100:14639–14644. 10.1073/pnas.2536769100

54. Luecke H, Schobert B, Lanyi JK, Spudich EN, Spudich JL (2001) Crystal structure of sensory rhodopsin II at 2.4 angstroms: insights into color tuning and transducer interaction. Science 293:1499–1503. 10.1126/science.1062977

55. Martin J, Bruno VM, Fang Z, Meng X, Blow M, Zhang T, Sherlock G, Snyder M, Wang Z (2010) Rnnotator: an automated de novo transcriptome assembly pipeline from stranded RNA-Seq reads. BMC Genomics 11:663. 10.1186/1471-2164-11-663

56. McGuffin LJ, Bryson K, Jones DT (2000) The PSIPRED protein structure prediction server. Bioinformatics 16:404–405. 10.1093/bioinformatics/16.4.404

57. Miranda MZ, Lichner Z, Szaszi K, Kapus A (2021) MRTF: Basic Biology and Role in Kidney Disease. Int J Mol Sci 2210.3390/ijms22116040

58. Mouilleron S, Wiezlak M, O’Reilly N, Treisman R, McDonald NQ (2012) Structures of the Phactr1 RPEL domain and RPEL motif complexes with G-actin reveal the molecular basis for actin binding cooperativity. Structure 20:1960–1970. 10.1016/j.str.2012.08.031

59. Nack M, Radu I, Gossing M, Bamann C, Bamberg E, von Mollard GF, Heberle J (2010) The DC gate in Channelrhodopsin-2: crucial hydrogen bonding interaction between C128 and D156. Photochem Photobiol Sci 9:194–198. 10.1039/b9pp00157c

60. Nagata T, Inoue K (2021) Rhodopsins at a glance. J Cell Sci 13410.1242/jcs.258989

61. Nagel G, Szellas T, Huhn W, Kateriya S, Adeishvili N, Berthold P, Ollig D, Hegemann P, Bamberg E (2003) Channelrhodopsin-2, a directly light-gated cation-selective membrane channel. Proc Natl Acad Sci U S A 100:13940–13945. 10.1073/pnas.1936192100

62. Panzer S, Zhang C, Konte T, Brauer C, Diemar A, Yogendran P, Yu-Strzelczyk J, Nagel G, Gao S, Terpitz U (2021) Modified Rhodopsins From *Aureobasidium pullulans* Excel With Very High Proton-Transport Rates. Front Mol Biosci 8:750528. 10.3389/fmolb.2021.750528

63. Pettersen EF, Goddard TD, Huang CC, Couch GS, Greenblatt DM, Meng EC, Ferrin TE (2004) UCSF Chimera--a visualization system for exploratory research and analysis. J Comput Chem 25:1605–1612. 10.1002/jcc.20084

64. Prasongsuk S, Lotrakul P, Ali I, Bankeeree W, Punnapayak H (2018) The current status of *Aureobasidium pullulans* in biotechnology. Folia Microbiol (Praha) 63:129–140. 10.1007/s12223-017-0561-4

65. Purschwitz J, Muller S, Kastner C, Fischer R (2006) Seeing the rainbow: light sensing in fungi. Curr Opin Microbiol 9:566–571. 10.1016/j.mib.2006.10.011

66. Rao F, Xue T (2024) Circadian-independent light regulation of mammalian metabolism. Nat Metab 6:1000–1007. 10.1038/s42255-024-01051-6

67. Rodaki A, Bohovych IM, Enjalbert B, Young T, Odds FC, Gow NA, Brown AJ (2009) Glucose promotes stress resistance in the fungal pathogen *Candida albicans*. Mol Biol Cell 20:4845–4855. 10.1091/mbc.e09-01-0002

68. Saito R, Smoot ME, Ono K, Ruscheinski J, Wang PL, Lotia S, Pico AR, Bader GD, Ideker T (2012) A travel guide to Cytoscape plugins. Nat Methods 9:1069–1076. 10.1038/nmeth.2212

69. Salentin S, Schreiber S, Haupt VJ, Adasme MF, Schroeder M (2015) PLIP: fully automated protein-ligand interaction profiler. Nucleic Acids Res 43:W443–447. 10.1093/nar/gkv315

70. Sancar A (2004) Photolyase and cryptochrome blue-light photoreceptors. Adv Protein Chem 69:73–100. 10.1016/S0065-3233(04)69003-6

71. Scheib U, Stehfest K, Gee CE, Korschen HG, Fudim R, Oertner TG, Hegemann P (2015) The rhodopsin-guanylyl cyclase of the aquatic fungus *Blastocladiella emersonii* enables fast optical control of cGMP signaling. Sci Signal 8:rs8. 10.1126/scisignal.aab0611

72. Shannon P, Markiel A, Ozier O, Baliga NS, Wang JT, Ramage D, Amin N, Schwikowski B, Ideker T (2003) Cytoscape: a software environment for integrated models of biomolecular interaction networks. Genome Res 13:2498–2504. 10.1101/gr.1239303

73. Shimono K, Ikeura Y, Sudo Y, Iwamoto M, Kamo N (2001) Environment around the chromophore in pharaonis phoborhodopsin: mutation analysis of the retinal binding site. Biochim Biophys Acta 1515:92–100. 10.1016/s0005-2736(01)00394-7

74. Sineshchekov OA, Jung KH, Spudich JL (2002) Two rhodopsins mediate phototaxis to low- and high-intensity light in *Chlamydomonas reinhardtii*. Proc Natl Acad Sci U S A 99:8689–8694. 10.1073/pnas.122243399

75. Tamura K, Stecher G, Kumar S (2021) MEGA11: Molecular Evolutionary Genetics Analysis Version 11. Mol Biol Evol 38:3022–3027. 10.1093/molbev/msab120

76. Trott O, Olson AJ (2010) AutoDock Vina: improving the speed and accuracy of docking with a new scoring function, efficient optimization, and multithreading. J Comput Chem 31:455–461. 10.1002/jcc.21334

77. Vaishnav P, Demain AL (2011) Unexpected applications of secondary metabolites. Biotechnol Adv 29:223–229. 10.1016/j.biotechadv.2010.11.006

78. Verchere A, Ou WL, Ploier B, Morizumi T, Goren MA, Butikofer P, Ernst OP, Khelashvili G, Menon AK (2017) Light-independent phospholipid scramblase activity of bacteriorhodopsin from *Halobacterium salinarum*. Sci Rep 7:9522. 10.1038/s41598-017-09835-5

79. Vickerman BM, Zywot EM, Tarrant TK, Lawrence DS (2021) Taking phototherapeutics from concept to clinical launch. Nat Rev Chem 5:816–834. 10.1038/s41570-021-00326-w

80. von Mering C, Huynen M, Jaeggi D, Schmidt S, Bork P, Snel B (2003) STRING: a database of predicted functional associations between proteins. Nucleic Acids Res 31:258–261. 10.1093/nar/gkg034

81. Wang J, Chitsaz F, Derbyshire MK, Gonzales NR, Gwadz M, Lu S, Marchler GH, Song JS, Thanki N, Yamashita RA, Yang M, Zhang D, Zheng C, Lanczycki CJ, Marchler-Bauer A (2023) The conserved domain database in 2023. Nucleic Acids Res 51:D384–D388. 10.1093/nar/gkac1096

82. Waschuk SA, Bezerra AG, Jr., Shi L, Brown LS (2005) *Leptosphaeria* rhodopsin: bacteriorhodopsin-like proton pump from a eukaryote. Proc Natl Acad Sci U S A 102:6879–6883. 10.1073/pnas.0409659102

83. Watanabe HC, Welke K, Schneider F, Tsunoda S, Zhang F, Deisseroth K, Hegemann P, Elstner M (2012) Structural model of channelrhodopsin. J Biol Chem 287:7456–7466. 10.1074/jbc.M111.320309

84. Waterhouse A, Bertoni M, Bienert S, Studer G, Tauriello G, Gumienny R, Heer FT, de Beer TAP, Rempfer C, Bordoli L, Lepore R, Schwede T (2018) SWISS-MODEL: homology modelling of protein structures and complexes. Nucleic Acids Res 46:W296–W303. 10.1093/nar/gky427

85. Yadav AK, Polasek-Sedlackova H (2024) Quantity and quality of minichromosome maintenance protein complexes couple replication licensing to genome integrity. Commun Biol 7. 10.1038/s42003-024-05855-w

86. Yang X, Jiang X, Yan W, Huang Q, Sun H, Zhang X, Zhang Z, Ye W, Wu Y, Govers F, Liang Y (2021) The Mevalonate Pathway Is Important for Growth, Spore Production, and the Virulence of *Phytophthora sojae*. Front Microbiol 12:772994. 10.3389/fmicb.2021.772994

87. Yoshida K, Tsunoda SP, Brown LS, Kandori H (2017) A unique choanoflagellate enzyme rhodopsin exhibits light-dependent cyclic nucleotide phosphodiesterase activity. J Biol Chem 292:7531–7541. 10.1074/jbc.M117.775569

