## Supplementary material for "*In silico* characterization of unique fungal modular rhodopsin expands the horizon of novel optobiological and biomedical applications": Fig S1, Fig S2, Fig S3, Fig S4, Fig S5, Fig S6, Fig S7

**Supplementary Figure 1:**

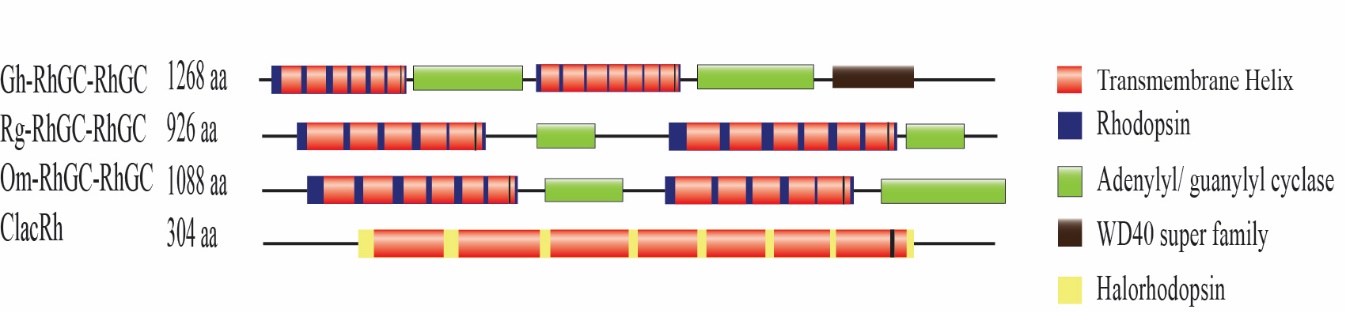

**Fig. S1** The schematic illustration depicts novel fungal twin rhodopsins domain and halorhodopsin. Twin rhodopsin guanylate cyclase (RhGC-RhGC) in *Gorgonomyces haynaldii* (Gh-RhGC-RhGC)*, Rhizoclosmastium globosum* (Rg-RhGC-RhGC) and *Obelidium mucromatum* (Om-RhGC-RhGC). The ClacRh from *Cladophialophora carrionii* represents the fungal halorhodopsin identified in this study. The blackline denotes the full-length protein, and domains are illustrated by geometric structures.

**Supplementary Figure 2:**

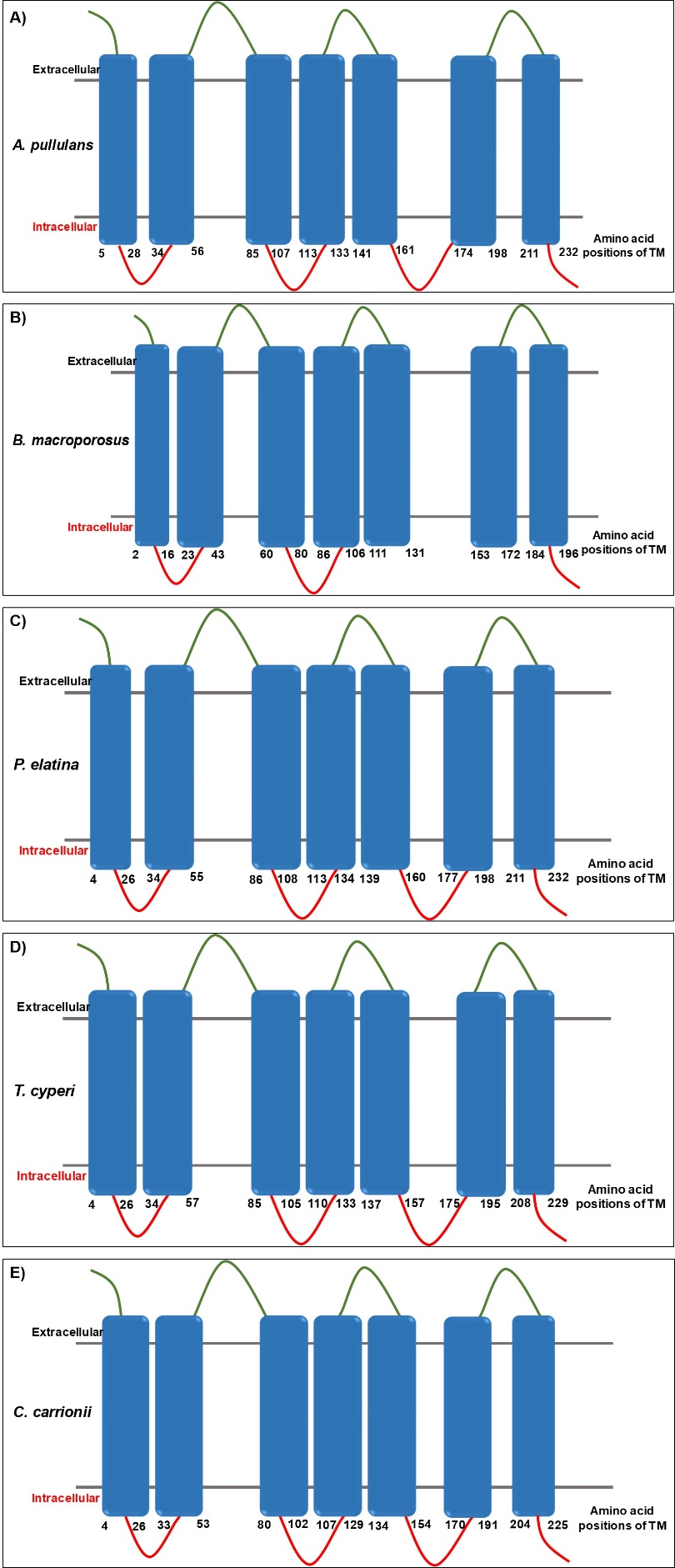

**Fig. S2** Schematic representation of the seven transmembrane helices (7TM) of the identified modular rhodopsins and halorhodopsin in the fungal genome predicted using DeepTMHMM-1.0 server. **A)** *Aureobasidium pullulans* (Rh-RPEL). **B)** *Boothiomyces macroporosus* (Rh-GC-cAT). **C)** *Pseudographis elatina* (Rh-NAD(P)). **D)** *Testicularia cyperi* (Rh-MCM). **E)** *Cladophialophora carrionii* (halorhodopsin). Blue represents the transmembrane helices, green represents extracellular loop and red represents the intracellular loop.

**Supplementary Figure 3:**

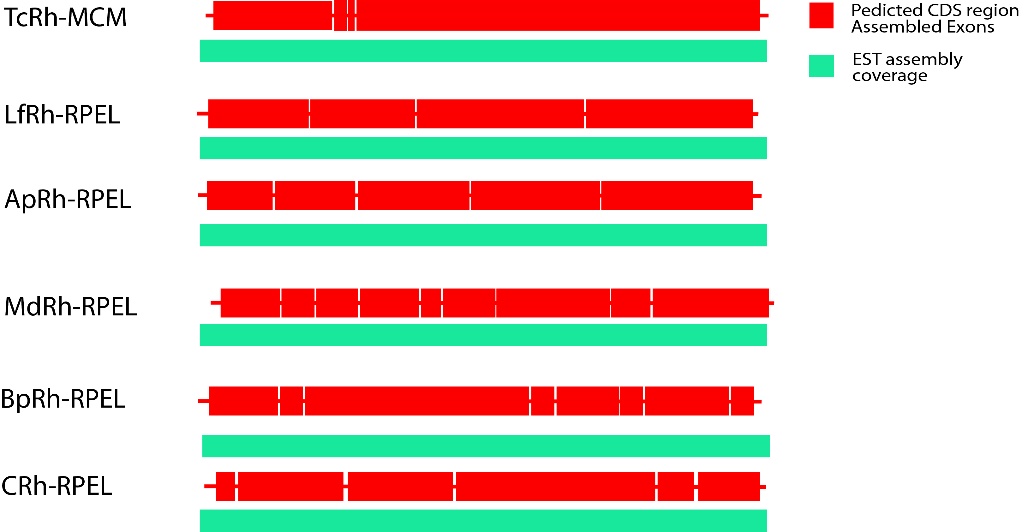

**Fig. S3** Experimental evidence of single open reading frame (ORF)-encoding entire novel multi-domain fungal rhodopsins.

**Supplementary Figure 4:**

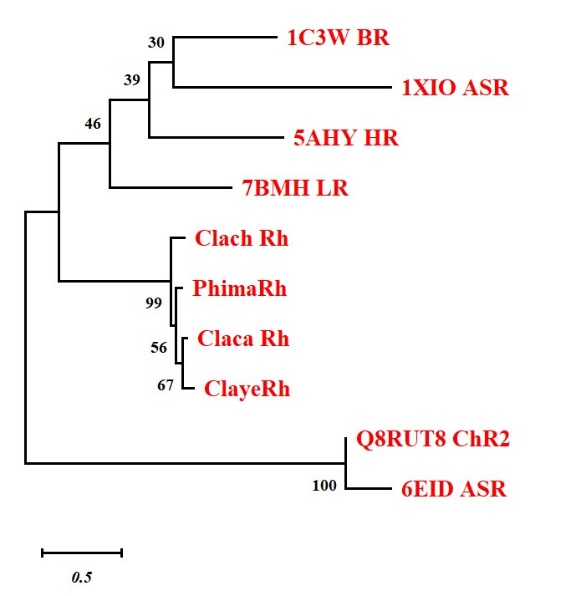

**Fig. S4** Maximum-likelihood tree showing the evolutionary positions of newly identified fungal rhodopsins (ClachRh, PhimaRh, ClacaRh, ClayeRh) relative to canonical microbial rhodopsins, including BR (1C3W), ASR (1XIO, 6EID), HR (5AHY), LR (7BMH), and ChR2 (Q8RUT8). Fungal sequences form a distinct, well-supported clade separate from classical Type I rhodopsins. Bootstrap values are shown at the nodes; scale bar indicates substitutions per site. (*Cladophialophora carrionii, Phialophora macrospora, Cladophialophora chaetospira , Cladophialophora yegresii CBS 114405*).

**Table S1** Structure validation by ERRAT and PROCHECK

| **Rhodopsin** | **pTM** | **ERRAT** | **PROCHECK** |
| --- | --- | --- | --- |
| TcRh_MCM | 0.93 | 100 | Out of 8 evaluations  Errors: 0  Warning: 4 ; Pass: 4 |
| PeRh_NAD(P) | 0.92 | 97.65 | Out of 8 evaluations  Errors: 0  Warning: 4 ; Pass: 4 |
| LfRh-RPEL | 0.93 | 100 | Out of 8 evaluations  Errors: 0  Warning: 3 ; Pass: 5 |
| ApRh-RPEL | 0.93 | 99.09 | Out of 8 evaluations  Errors: 0  Warning: 4 ; Pass: 4 |
| MdRh-RPEL | 0.93 | 99.06 | Out of 8 evaluations  Errors: 0  Warning: 3 ; Pass: 5 |
| BpRh_RPEL | 0.93 | 100 | Out of 8 evaluations  Errors: 0  Warning: 3 ; Pass: 5 |
| C.sp.Rh-RPEL | 0.93 | 99.54 | Out of 14 evaluations  Errors: 0  Warning: 5 ; Pass: 9 |
| EeRh_RPEL | 0.93 | 99.54 | Out of 8 evaluations  Errors: 0  Warning: 4 ; Pass: 4 |
| BmRh-GC-cAT | 0.9 | 97.56 | Out of 8 evaluations  Errors: 0  Warning: 2 ; Pass: 6 |
| TsRh-GC-cAT | 0.82 | 98.36 | Out of 8 evaluations  Errors: 0  Warning: 2 ; Pass: 6 |

**Supplementary Figure 5:**

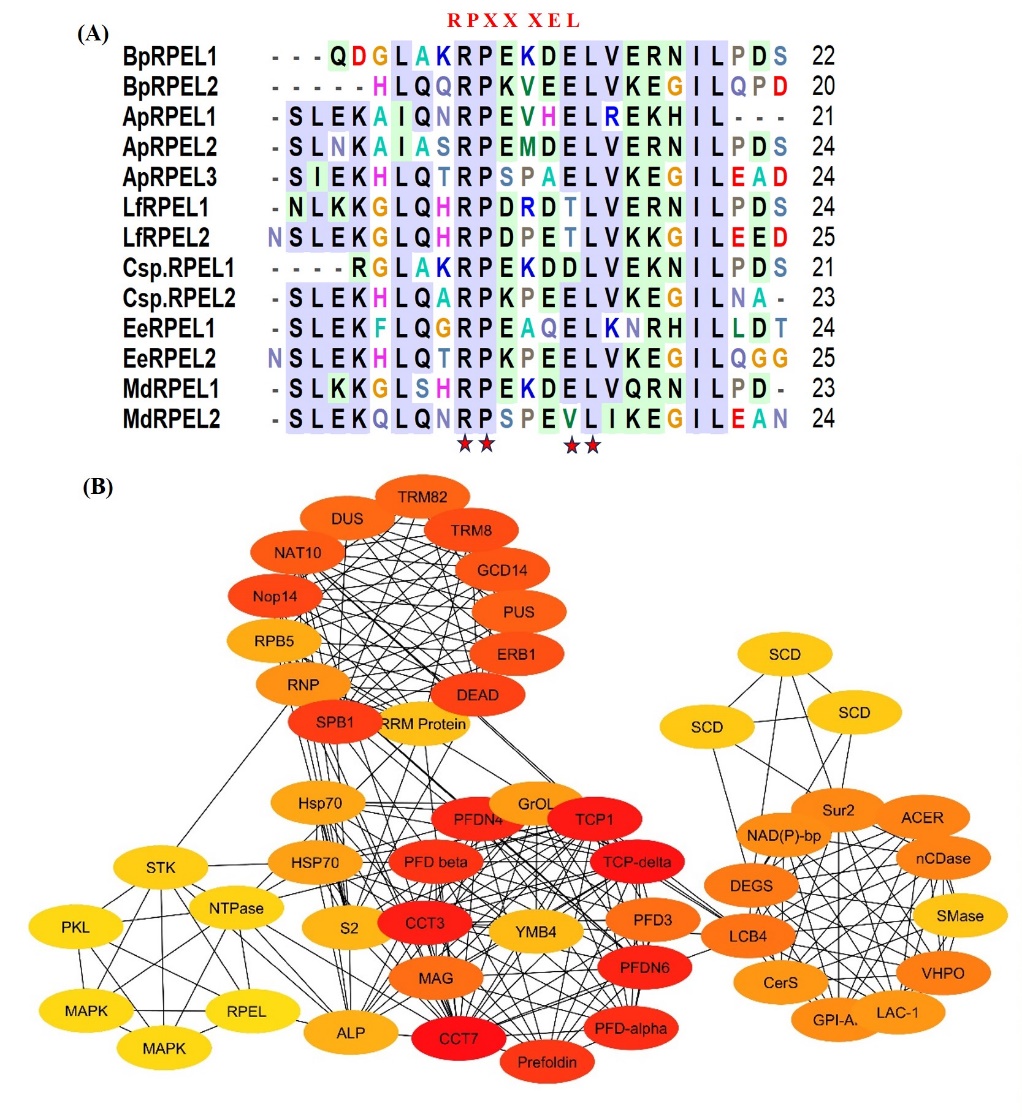

**Fig. S5 (A)** Sequence alignment of individual RPEL motifs of identified modular rhodopsin. Conserved motifs are highlighted by an asterisk (*). **(B)** Protein-Protein interaction network showing interacting partners of RPEL repeats domains in *Aureobasidium pullulans*. Protein-protein interaction was performed using String version 11 (<https://string-db.org/>). The principal nodes that control the overall network were analysed by CytoHubba analysis using the betweenness-centrality algorithm. Colour range (orange red to yellowish) indicates scores ranked by betweenness method.

**Supplementary Figure 6:**

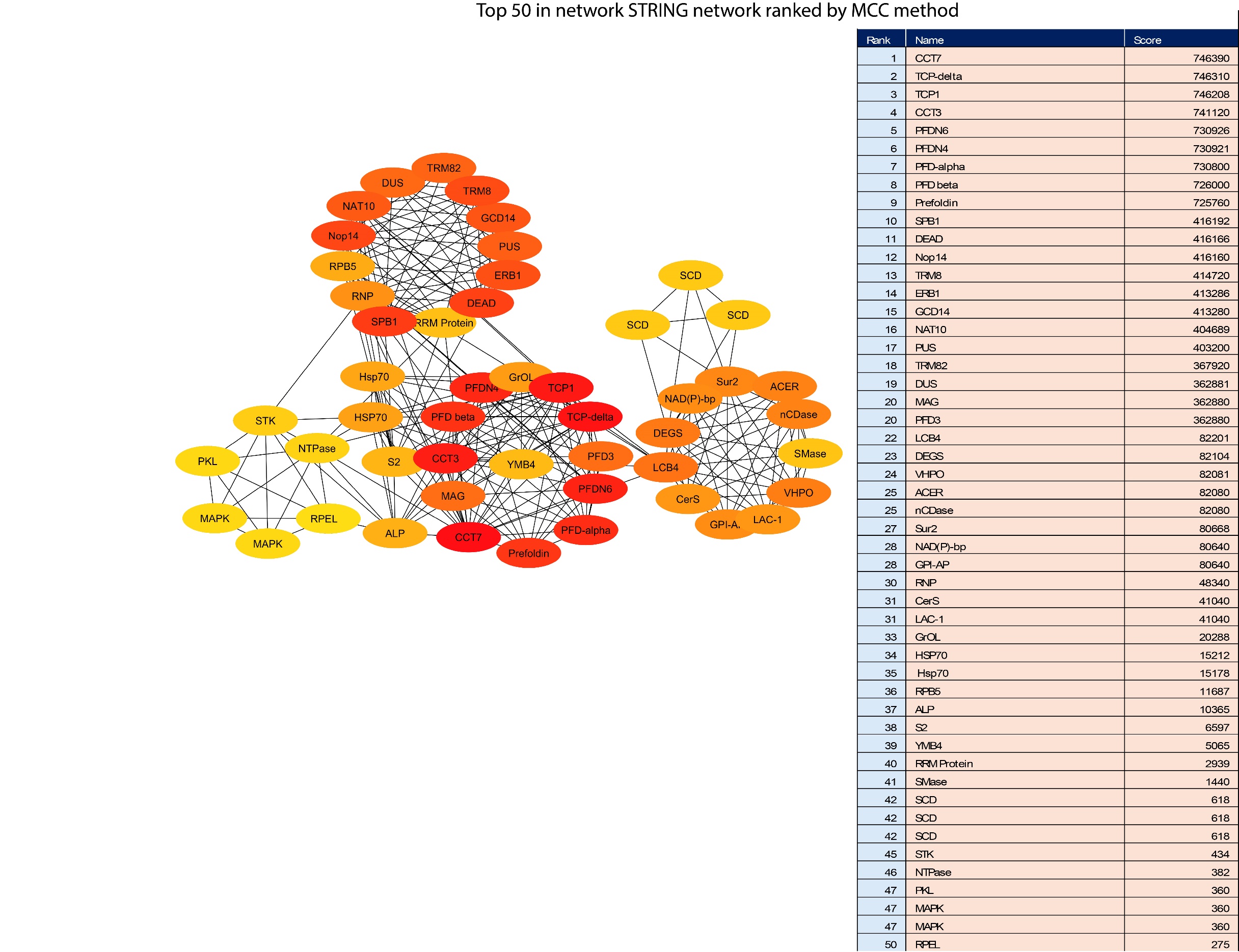

**Fig. S6** Top 50 predicted interacting partners involved in the crosstalk between RPEL effector domain in *Aureobasidium pullulans* with protein modification and sphingolipid metabolism pathways.

**Supplementary Figure 7:**

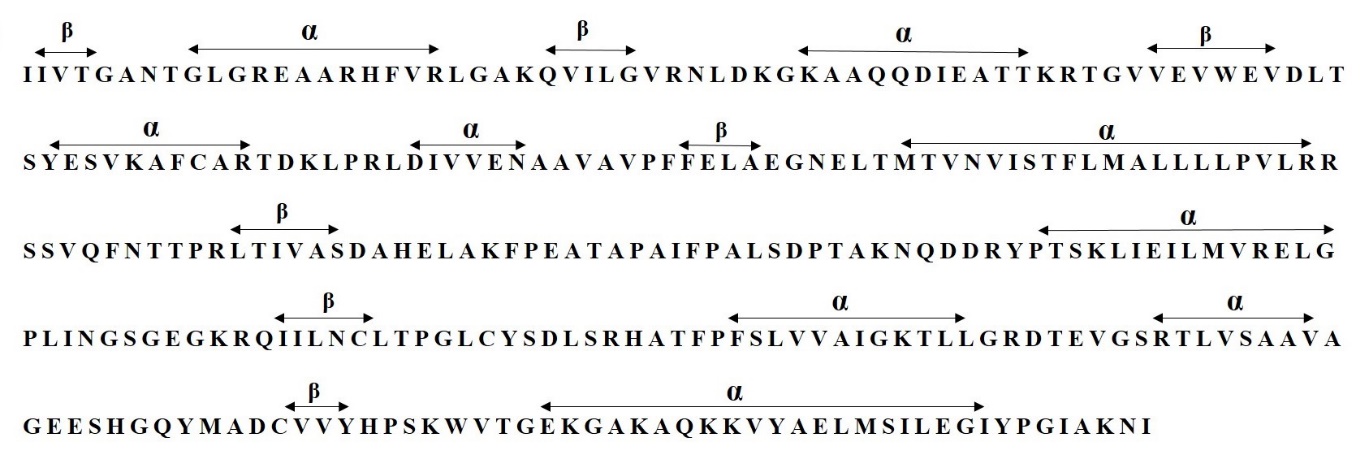

**Fig. S7** Sequence of NAD(P) Rossmann fold in *Pseudographis elatina* with marked α-helices and β-strands.

**Protein sequences of the effector domains used in the current study:**

The following table denotes the sequences that have been employed for the study of modularity, homology, phylogenetic tree, structural and Biosynthetic Gene Cluster (BGC) analysis. The full-length sequences are also appended.

| ***Phylogenetic pattern of newly identified modular fungal rhodopsins*** | | |
| --- | --- | --- |
| **Organism** | **Domain** | **Sequence (amino acids)** |
| *Capnodiales* sp. | Rhodopsin | 39-271 |
| *Elasticomyces elasticus* | Rhodopsin | 42-274 |
| *Baudoinia panamericana* | Rhodopsin | 39-271 |
| *Aureobasidium pullulans* | Rhodopsin | 89-321 |
| *Myriangium duriaei* | Rhodopsin | 37-269 |
| *Lentithecium fluviatile* | Rhodopsin | 46-276 |
| *Pseudographis elatina* | Rhodopsin | 66-298 |
| *Terramyces* | Rhodopsin | 65-271 |
| *Cladophialophora carrionii* | Rhodopsin | 24-249 |
| *Halobacterium salinarum* | Bacteriorhodopsin | 1-231 |
| *Anabaena* | *Anabaena* Sensory Rhodopsin (ASR) | 1-261 |
| *Chlamydomonas reinhardtii* | Channelrhodopsin | 1-737 |
| *Boothiomyces macroporosus* | Rhodopsin | 65-271 |
| *Testicularia cyperi* | Rhodopsin | 29-258 |
| ***Homology analysis of the microbial rhodopsins*** | | |
| *Anabaena* | *Anabaena* Sensory Rhodopsin (ASR) | 1-261 |
| *Halobacterium salinarum* | Bacteriorhodopsin | 1-231 |
| *Pseudographis elatina* | Rhodopsin | 66-298 |
| *Lentithecium fluviatile* | Rhodopsin | 46-276 |
| *Myriangium duriaei* | Rhodopsin | 37-269 |
| *Aureobasidium pullulans* | Rhodopsin | 89-321 |
| *Baudoinia panamericana* | Rhodopsin | 39-271 |
| *Capnodiales* sp. | Rhodopsin | 39-271 |
| *Elasticomyces elasticus* | Rhodopsin | 42-274 |
| *Testicularia cyperi* | Rhodopsin | 29-258 |
| *Cladophialophora carrionii* | Rhodopsin | 24-249 |
| ***Structural analysis of microbial rhodopsin*** | | |
| *Aureobasidium pullulans* | Rhodopsin | 89-321 |
| *Anabaena* | *Anabaena* Sensory Rhodopsin (ASR) | 1-261 |
| *Baudoinia panamericana* | Rhodopsin | 39-271 |
| *Lentithecium fluviatile* | Rhodopsin | 46-276 |
| *Myriangium duriaei* | Rhodopsin | 37-269 |
| *Capnodiales* sp. | Rhodopsin | 39-271 |
| *Elasticomyces elasticus* | Rhodopsin | 42-274 |
| *Leptosphaeria maculans* | Light-gated proton rhodopsin | 49-281 |
| ***Domain analysis of the newly identified modular fungal rhodopsins*** | | |
| *Aureobasidium pullulans* | Rh-RPEL | FL: 1-552  Rh: 89-321  RPEL (1): 434-454  RPEL (2): 478-501  RPEL (3): 522-545 |
| *Baudoinia panamericana* | Rh-RPEL | FL: 1-376  Rh: 39-271  RPEL (1): 305-326  RPEL (2): 347-370 |
| *Lentithecium fluviatile* | Rh-RPEL | FL: 1-382  Rh: 46-276  RPEL (1): 308-331  RPEL (2): 352-375 |
| *Myriangium duriaei* | Rh-RPEL | FL: 1-467  Rh: 37-269  RPEL (1): 349-372  RPEL (2): 393-416  RPEL (3): 437-460 |
| *Capnodiales* sp. | Rh-RPEL | FL: 1-367  Rh: 39-271  RPEL (1): 296-316  RPEL (2): 337-360 |
| *Testicularia cyperi* | Rh-MCM | FL: 1-1401  Rh: 29-258  MCM (1): 446-565  PLPDE: 610-891  MCM (2): 902-1377 |
| *Boothiomyces macroporosus* | Rh-GC-cAT | FL: 1-1154  Rh: 65-271  AC: 318-492  cAT: 516-1095 |
| *Rhizoclosmastium globosum* | RhGC-RhGC | FL: 1-929  Rh (1): 116-258  GC (1): 284-465  Rh (2): 544-746  GC (2): 772-921 |
| *Gorgonomyces haynaldii* | RhGC-RhGC | FL: 1-1267  Rh (1): 63-238  GC (1): 260-443  GC (2): 701-892  WD40: 902-1038  Rh (2): 473-681 |
| *Obelidium mucromatum* | RhGC-RhGC | FL: 1-1088  Rh (1): 133-317  GC (1): 343-524  Rh (2): 713-856  GC (2): 907-1087 |
| *Pseudographis elatina* | Rh-NAD(P) Rossmann | FL: 1-672  Rh: 66-298  NADB Rossmann: 372-639 |
| ***Biosynthetic Gene Cluster (BGC) analysis*** | | |
| *Aureobasidium pullulans* | Rh-RPEL | FL: 1-552  Rh: 89-321  RPEL (1): 434-454  RPEL (2): 478-501  RPEL (3): 522-545 |
| FL: Full length sequence | | |

**The full-length sequences are as follows:**

**>****TIA65092.1 family A G protein-coupled receptor-like protein [*Aureobasidium pullulans*] (Rh-RPEL)**

MTTSSRQAGAIESSARSSSELAAALFKNVSPSAALPSHPQPLSSLHPIPSSIHAPAMIVDPVEAFKATSSVAPIPTVVPSLPEYETVTETGTRALWAVFVLMLLSMIVFVGLSWTVPISKRLYHVVTTLIVTFASLSYFAMATGHGISYHRTTVTDSHRHVPDTTHDVYRQVYWARYVDWSLTTPLLLLDLALLAGLSGGHILLAIVADVIMVLTGLFAAYGTEGTPQKWGWYAIACIAYLVVIWMLAVHGRANAMAKGGKVGKFFASIAGFTLVIWTIYPIVWGVADGSRKMSVDQEIIAYAVLDLLAKPVFGAWLLFTHQSMPETQVELGGFWTHGVSSEGAIRVDPVTTPVNDNPARFMQGLSLSGEEGAQELTHHLNSHLPAHNDICFCGSTRLPKLLYCSGTQQQAPATMSDELTPATTSPISLERRNSLEKAIQNRPEVHELREKHILLNTNAAPALQAQQQELQRHKLTDSLNKAIASRPEMDELVERNILPDSTAAPALQNHQRELAAAMRRDSIEKHLQTRPSPAELVKEGILEADENPLDGP

**>XP_007678484.1 hypothetical protein BAUCODRAFT_149869 [*****Baudoinia panamericana* UAMH 10762] (Rh-RPEL)**

MIVDPVQAFATGTIPIATSSATPLPTIIPSLPEYQTATDTGNRTLWVVFVVMLVASIVFAGLSWNVPMSKRLYHIVTCLITIFASLSYFAMATGHGIGYHHVVERESHKHVPDTTYDVYREVYWARYVDWSLTTPLLLLDLCLLAGISGGNIMIAIVADIIMILGGLFAAFGSEGTPQKWGWYTIACIAYLVVIWQLAYNGRAMAMSKGGKVGNFFAAIGGFTLVIWTVYPIIWGIADGSRNMNVDEEIIAYAVLDILAKPVFGTWLIYTHMTMPETNVEIGGFWSEGLKGEGQLRVGDDEGKQDGLAKRPEKDELVERNILPDSMAAPALQEKQRELEKHMRADSLEKHLQQRPKVEELVKEGILQPDENPIAEG

**>****KAF2681323.1 family A G protein-coupled receptor-like protein [*****Lentithecium fluviatile* CBS 122367] (Rh-RPEL)**

MIIDPVEILKKTSTAGPIPTATPTSVDPIPTVLPDSPEKQFVGDGGTRTLWVVFIVMLISSAVFAGLSWRVPVGRRLYHVITTLITIFAAISYFAMATGHGVSVHTIQVRHQIDHLPDTFTEVQRQVFWARYVDWSLTTPLLLLDLSLLAGLNGAHILMAIVADIIMILTGLFAAFGSEGTPQKWGWYAIACIAYLVVIWHLAVNGRAQAQAKGDKVGSFFLAIAGFTLIVWTAYPIVWGIADGSRNLSVDGEIIAYAVLDILAKPVFGTWLLIAHARMPETNIDLGGFWSYGLGGEGSVRLGDDDDNLKKGLQHRPDRDTLVERNILPDSNAAPALQGHQKELERHMRANSLEKGLQHRPDPETLVKKGILEEDENPLKDA

**>jgi|Myrdu1|313850|estExt_fgenesh1_pg.C_7_t10474 [*Myriangium duriaei*] (Rh-RPEL)**

MIVDPNQVAAFAAAKTSVAPVPSFTPGPSVSYETAGDSGKRALWVVFVIMLVATVVFSFLSFSVPISKRLYHVITTLIVTFAALSYFAMASGDGISLHKNVVTEEHKHVPDTQTYVYREVYWARYVDWSLTTPLLLLDLALLAGLSGGNIVIAVVADIIMVLTGLFAAFGKEESPSKWGWYAIACIAYLVIVWQLAVNGRATAFGKGGKVGTFFASIAGFTLIVWTIYPIVWGVADGARIASVDQEIIAYAVLDVLAKPVFGAWLLYTHAAIPETNIEVGGFWTHGVSGEGAIRSVAVVEFFPDLVPSLVERTHFNTLTTLMTMAANSDTTPAIGVDDSPINTLERRNSLEKHLQTRPEEQDLKNRHILLDTNAAPALQAQQHELERQRITDSLKKGLSHRPEKDELVQRNILPDSNAAPALQGKERELSKHMRANSLEKQLQNRPSPEVLIKEGILEANENPLADS

**>****jgi|Cap6580_1|992589|estExt_fgenesh1_pm.C_720012 [*Capnodiales* sp.] (Rh-RPEL)**

MLVDPVKAFATGILPLPTASASPLPSVVPSAAEYQDATETGNKTLWVVFTIMFIASIVFSGMSWNVPISKRLYHIVTTMIVIFASLSYFAMASGHGISYHHIEIRESHKHVPDTTKDIYRQVYWARYVDWSLTTPLLLLDLCLLAGLSGGHILLTMVADIIMILTGLFAAFGTEDTPQKWGWYAIACVAYLVVIWHLVIHGRATAMSKGGKVGNFFAAIGGFTLVIWTVYPIIWGIADGSRNMNVDEEIIAYAVLDVLAKPIFGAWLLFTHMSMPETDVDLGGFWSEGLKGEGNIRGLAKRPEKDDLVEKNILPDSTAAPALQERRRELERHMRADSLEKHLQARPKPEELVKEGILNANENPVAES

**>****KAK4972783.1 hypothetical protein LTR42_006077 [*****Elasticomyces elasticus*] (Rh-RPEL)**

MIDPVQAFATAMASATLPLATSSVSPIPTVVPTLPEYQDATETGTRTLWVVFVIMLLATVVFSGMAWTVPISKRLYHIVTTLIVIFASLSYFAMASGHGISYHHVVVRESHKHVPDTTTDLYRQVYWARYVDWSLTTPLLLLDLALLAGLSGGVILMIMVADIIMILTGLFAAFGTEDTPQKWGWYAIACIAYLVVIWHLVIHGRATAMNKGGKVGNFFAAIGGFTLVIWTVYPIIWGIADGSRNMNVDEEIIAYAVLDILAKPVFGAWLLFTHASMPESNVELGGFWSEGLKGEGNIRVGDDDEATKQQYEIMAEQEANNAVDRSPIAQLERRNSLEKFLQGRRPEAQELKNRHILLDTTSAPTLQARQAELERQRITDSLKRGLAKRPEKGDLVEKNILPESTAAPSLQETQKKLAKQMRADSIEKHLQGRPKQEGILHADETPIAEG

**>****KAJ3272524.1 hypothetical protein HDV01_005475** **[*Terramyces* sp. JEL0728] (Rh-GC-cAT)**

MNQNDLYLFDRGNSDGWYKNNASAFVISGTFAFFANLTANSPDKKSLAVVLLLVNLITLCSYLLMAWRLTPQFKDGNQYPVDVARYLEWIATCPNLILLVGEFTKENTFTFRIFVYDYLMLISGFLASTTTGSISTISLLMSCIYFVLIMQGLHFMFTKAIAGGTLCVLDKQTLQIAHASTMVGWTIYIIAKAFLTIVLVNATIEQTQSETVQNISGIVATMEKEFNNTEAILEKVFPSELLPEIKEGNFPEAKDYKSVTIFFSDITNFNSITSKSSTRDVLKTLDALWTKYEEIGRKWGICKVETIGDAFLGISGCPTRTDNHAENCLNFALEIIEMAKEFKTSANDSIHIRIGIHSGPVVAGFMGTNNPRWCILGDAVNTASRIETLSKPMKINISESTFELVKEKNLFEFSEPHIYEIRGKGRFNMYWVAAKPRSINRNLVAKRFQQTFASQGKVPRLPIPTLENLEQKYLKSCQPLLAKEDYEKTEAIVKEFLSKVDHPEQPSDLLTKPPPKGVLTSFQIKRAAGLITNLLNFNDMLNNQTFPVEAIKGTPLCMNQYKNIFGTTRLPGKTSDTLSHQYPTTAKHIIVLTKNEIYKVQVLADDGKRVSNAELERVLLNIGKETLVDRHPPHIGVLTAGDRDTWYEASEKLCALSPSNVKNFDIIKDSLFAVCLDDHSTKKNLDLSLTQIFHNNNAQNRWFDKSLQLIITNSGRAGLNGEHTPSDAVVPGNVMDYIISHEPSIDPQNTVQGQYLPPPQRLEWVVDDSVNSLIEKARGVAQALIDDTESTLLQTDYYGSRFMKEIAKTSPDAYIQIALQLAYYRLHQKPTAVYESASTRFFKHGRTETGRSMSNESLEFISTFDNDDVLYDTKRELFRKAIATQSNYMKDAAYGKGIDRHMLGLRCMIKPDEMEKATMFTDPAYITSMTFRLSSSNMSPGRNFYGGFGPVVFDGYGINYAIDKDNLKFSISAKRSCTETKIYRLRDELEKVFKDLCILFPKRSEIWGKNWRKEQQAEKVADIRFAKMKTLSDEYLSKQASLAEKYLDRK

**>KAJ3254479.1 hypothetical protein HK103_007115 [*Boothiomyces macroporosus*] (Rh-GC-cAT)**

MSNLWSAKIKNETEKSRIQSTTINQMILSLFITVPMAVYTHYVHSGAQPFDRGNADSWYNYNASIFTFTGLSAFFAMLTAKSNDKKSLAIVLGLVNLITLCSYLTIAKRLTPNVPDSNGYPVDFARYLEWIACCPNMVLLIGEFTKDAEYTSQTFLNNYVMLCCGFLASVTGESLSIIFTSITVYLFIKVMTGLHRMYSNAIDGRTGCTLDRTMVRIANITTTFGWSLFPIVWFLVRFKVISFETGEMLYCFGDIVAKAFLTLVLVNATIEHAQNETIQRISGIASVMEREMNNTDVVLEKFIPPEVLSNIKEGNFPEAQDYESVTILLSDITNYNVLSARNSTRDVLKTLDSLWSKYEEIGKKWGIYKVETIGDAYLGVSGCPTRSENHAENCLNFALDILEMTTEFKTETKEGIQIRIGIHSGPVVAGLMGANNPRWCIVGDSVSTASRIEALSKPMRINISESTYELVKDKNMFEFSGPNSYEVKGKGRSLNRNLLTKRFQQTFANQDKVPRLPIPTLENLEQKYLKSCQPLLSKEDYEKTESIVKEFLSKDGFGTTLQERLIEYEKKEPHSWLEKIWLQKAYLEWREPSMINVNWWCQLVDHPEHPLDLLTKPPPKGVLTSFQIKRAAGLITNLLNFNDMLNNQTFPAESIKGTPLCMNQYKNIFGTTRLPGKTSDSLSHQYPTTAKHIIVLTKNEIFKVQVLTDDGKRVPIAEIERVLLNIGKETLVERSPPHIGVLTAGDRDTWFEAYEKLRALSPTNVKNFDIIKDALFAVCLDDHSTKKNLDQSLTQIFHNNNAQNRWFDKSLQLIVANSGRAGLNGEHTPSDAVVPGNVMDYIISHEPSIDPENAAQGEYLAPPQRLDWVVDDSVNALIEKARGVAQALIDDTESTLLQTDYYGSRFMKEVAKTSPDAYIQIALQLAYYRLHKKPTAVYESASTRFFKHGRTETGRSMSNESLEFISTFDNDDVLYDSKRELFKKAVNTQSNYMKDAAYGKGIDRHMLGLRCMIKPDEMDKATMFTDPAYITSMTFRLSTSNMSPGRNFYGGFGPVVFDGYGINYAIDKDNLKFSISAKRSCTETKIYRFKDELEKVFKDLCILFPKRSEVWGKNWRKDQQAEKVADIRFAKMKALSDEYLTKQASLAEKYSKKA

**>jgi|Rhihy1|734731|fgenesh1_pg.8_#_202 [*Rhizoclosmastium globosum* ] (RhGC-RhGC)**

MAEKDNSGAVYYVFRNFVVWSVLTVSGFYLLGTGYRGPTFDRGIADEFYAYTAASFGIAVIYSYIAYANAQNNEKRNLGKVLCIVNFIAMSSYIMQYTRTTPAFTDYVGYPVDPSRFYEWLATCPILIYLIAEITDNHHMADTTASYDYSLIILGFIACFLKQPYSEWFACGSTIFFFYTIYGLMVMFNSAIAGRTGCKLDVNSLYYARFVTFLAWNSFATVWYIQRAQIVTYEQGEVLFCVSDIFAKVFLTFILVNATLEESMNSKAKKMEAVASEIESQMAQADKLLEKLMPPSIVEAMKAGKANGAEEYASVTVFFSDITNFVALSGKSSTKDMLATLNKLWVEYDVICKRWGVYKVETIGDAFLGVVGAPDRIPDHAERAANFAIDVIRMVQDFRTVNDEEIVTRIGLNSGPITAGILGDSNPHWCIVGDAVNTASRMESTSKPMRIHISENTYKLINGKGFKLEGPDVMNIKETDWKRKIEAAKAEKDQSGTVFYVYRNYAAWMVVTLFGFYQLANGHKGPTYERGFAIEAYAYTACCFGIAVIYSFIAFFNAQNNEKRDLGKVLCVVNFISMVSYLLQWTGYTVSYTDVYGHPTDPARFYEWISTCPILIYMIAEITDNHHLADNTASADYVLLVLGYSSTFLRQPFSEYAAISAATCFFFVVKGFSSMFDRAIEGKTNCKLDAGSLWNAKWVTILAWASFPLSFFTQRSGIVTYETGEMMFCVADIWSKVFLTFILVNATLEESMNSKAKKMEAVAHEIESQMAQADKLLEKLMPASIVEAMKAGKATGSEEYSSVTVFFSDVTNFHALSNKNSTKEMLATLNKLWIEYDVICKRWGVYKVETIGDAFLGVVGAPERVPDHAERAANFAVDVISMVQDFRTDKGEELITRVGLNSGPITAGILGDSNPHWCIVGDAVNTASR

>**jgi|Gorhay1|216295|fgenesh1 [*Gorgonomyces haynaldii*] (RhGC-RhGC)**

MSASKVSWKDKLDQVQHASKVSGSSISTIERNVWLWVPFTAVGAFYAAIGRKSRFDRSSADNWYYYTSSVFGIGIFFSLVALASAKSAEKRSLSQVLLWVNMIACSTYMLSATRLTYTFVDVNGYPVDVARYVEWISTCPVLILLIGEITKCGNIARTTMKYDYIMLILGFIGAITREPISFLFQMMCFAYFTLILSGLSTMYDLAIQGKTGANLDKASLITAKWATLLAWNCFTFVWFGVRYRVISFATEMESELSNSDALLQRMMPPEVIEQLKTGQAPGAEEYDSVTVFFSDITNFTVLSSQSSTKDMIGTLNKLWLEYDAIAKKWGVYKVETIGDAYLGVVGCPTRTPDHAAAAVNFALDIVEMVRGFKTEMGSSIQIRVGLNSGPITAGVLGDLNPHWCIVGDTVNTASRMESTSKPMHIHISESTYKLATKFGKFKITGPDVLQVKALPHRQKALRFDRGDGPFYDMWVASIFAFGALFALLAQGSARTAEKVALTKVLFMCELLSMSTYLIQAYRISVSLPSWNGWPVDVARFLEWITTCPILIQLISDVCKTPDLTHKTVTFDYILLVCGFMASITRNPYSFLFSWIAFGCFTQVVTGLWSMFQRAIDGNTACKLHPSALRQAQLATCVSWACFPTVWLLMQFKVVSYGTGELLYGIADIFAKVVLTLILVNATVEQAQSERVSALESIATDMEKELGNTGALLSRMMPPEVIEQLKSGRAPGAEEYDSVTVFFSDITNFTVLSSQTSTKDMLATLNKLWLEYDAIAKKWGVYKVETIGDAYLGVVGCPTRSPDHAANACSFALDIVEMVRGFKTEMGSSIQIRVGLNSGPITAGVLGDLNPHWCIVGDTVNTASRMESTSKPMHVHCSESTYKLANPSGRFRFAGPDVLQIKRGLLDAVNCLRGGHLGTVSSLVVVFESGLVDVYDLSEAQERLILSFDNRVSTWGIAIQSERQWIAVSSNSHFITLYDLKTGTQRQFQGHDHNIPGIEFMGDYLVSCSIDGTCRIWNGSDGQLLHTASHGEEWKWCVRHLSKTDAQVVDKHRIQRLKDDPIDVPLLERFQSDPVLQSDCETGSGFGSPDQRSVDSMESVQHSPLKRKSDLEEQIGSPKKQELSKIEEPPIPQLPTLRSLLEQDIPEDLEEMDEDWSQQNLSETESLEYESSIEEWVTLRPKGEISTNDLILLTSKNHVYLLDQELRLICGLSHHAQWDIRYGLERNLFDTRLIYGRVDSQEASASDSKSERPPDVCQARSAWVQHRS

**>jgi|Obemuc1|938368|gm1.9661_g [*Obelidium mucromatum*] (RhGC-RhGC)**

MDDIQRGNCDNCNCSVYQVQTGTRRCRGCNHGAIFHQVVVRVAPKTSGKTNWKDKLQAAMAEKDNSGAVYYVFRNFLVWSGLTGIGWYLLTTGYVAPRFDRGYADEFYAYTAACFGIAVIYSYIAYANAQNNEKRDLGKVLCIVNFIAMSSYLMQWTRTTPTFSDYVGYPVDPARFFEWLATCPILIYLIAEITDNHHMADLTASYDYTLIVLGFIACFLKQPYSEWMACCSTIFFYYTITGLMEMYSKAIDGKTGCKLDVASLKVAKVITFLAWNSFTITWYIQRAQIVTYEQGELIFCINDIFAKVFLTFILVNATLEESMNSKAKKMEAVAQEIENQMAQADKLLEKLMPPSIVEAMKAGKASGAEEYSSVTVFFSDITNFVQLSSNNTTKDMLANLNKLWVEYDVICKRWGMYKVETIGDAFLGVIGAPDRIPDHAERAANFAIDVIRMVQDFRTIKDEEIVTRIGLNSGPITAGILGDSNPHWCIVGDAVNTASRMESTSKPMKIHISENTYKLINGKGFKLEGPDVMNIKEEKAMDEIQRGTCDNCNCSVFQVQTGTKRCRGCHHGAIFHQVIVRAVPKSGKTNWKEKLEAAKAEKDNSGAVYYVFRNFIAWSVMTGLGWYLLATGYALSAFDRGYADEFYAYTAGCFAIAVIYSYIAYANAQNNEKRDLGKVLCIVNFIAMATYILQFTRTSPSFKDYVGYPVDPARFFEWIATCPILIYLIAEITDNHQMADQTATYDYVLITFGFFACFLKQPYSEWMACSAVIFFFYTIIGIVTMYTRAIEGKTSCKLDVPSLAAARAITFLAWSAFPVTWHLQRAQIVTYEQGEVMFCVSDIFAKVFLTFILVNATLEESMNSKAKKMESVAHEIEAQMAQADKLLEKLMPPSIVEAMKAGKASGAEEYSSVTVFFSDITNFVALSNKNSTKDMLANLNKLWIEYDVICKRWGMYKVETIGDAFLGVIGAPDRIPDHAERAANFAIDVIRMVQDFRTIKDEEIVTRIGLNSGPITAGILGDSNPHWCIVGDAVNTASRMESTSKPMRIHISENTYKLINGKGFKIEGPDVMNIKGKGTMNTYWLNGR

**>****jgi|Pseel1|2270|g191.t1 [*Pseudographis elatina*] (Rh-NAD(P) Rossmann)**

MRDPMEALMTMAVKTSSSSLLVPTSSLSIPTSSMTSSLLIPTSTPTSVAPIPTVKPDLPVYETVADAGTTTLWIVFVIMLFSTVAFATIAYKTPVQKRLFPVLTTFITLFATLSYFAMATGDGNSFTHIILRETHKGDIPTTVKHVFRQVFWARYVDWSVTTPLVLLDLSLLAGLNGANIIVAVVADVFMVLTGLFAAFGHSDGQKWGYYAMACIAYLSIIYLLAVPGRKAVSAKSKPTRTLFVSIALYTLVLWTLYPIIWGFGDGSRILSVDSEIIAYAVLDVLAKPGFGIWLLVAHSRIGSTTIEGFWAHGLAAEGIIRIDDDEGEGGWLVEGLQALEVVGPTRPFRIDFVKSQFKKLPYPTADFESQCIIVTGANTGLGREAARHFVRLGAKQVILGVRNLDKGKAAQQDIEATTKRTGVVEVWEVDLTSYESVKAFCARTDKLPRLDIVVENAAVAVPFFELAEGNELTMTVNVISTFLMALLLLPVLRRSSVQFNTTPRLTIVASDAHELAKFPEATAPAIFPALSDPTAKNQDDRYPTSKLIEILMVRELGPLINGSGEGKRQIILNCLTPGLCYSDLSRHATFPFSLVVAIGKTLLGRDTEVGSRTLVSAAVAGEESHGQYMADCVVYHPSKWVTGEKGAKAQKKVYAELMSILEGIYPGIAKNI

**>Q8RUT8_ChR2 [*Chlamydomonas reinhardtii*] Channelrhodopsin**

MDYGGALSAVGRELLFVTNPVVVNGSVLVPEDQCYCAGWIESRGTNGAQTASNVLQWLAAGFSILLLMFYAYQTWKSTCGWEEIYVCAIEMVKVILEFFFEFKNPSMLYLATGHRVQWLRYAEWLLTCPVILIHLSNLTGLSNDYSRRTMGLLVSDIGTIVWGATSAMATGYVKVIFFCLGLCYGANTFFHAAKAYIEGYHTVPKGRCRQVVTGMAWLFFVSWGMFPILFILGPEGFGVLSVYGSTVGHTIIDLMSKNCWGLLGHYLRVLIHEHILIHGDIRKTTKLNIGGTEIEVETLVEDEAEAGAVNKGTGKYASRESFLVMRDKMKEKGIDVRASLDNSKEVEQEQAARAAMMMMNGNGMGMGMGMNGMNGMGGMNGMAGGAKPGLELTPQLQPGRVILAVPDISMVDFFREQFAQLSVTYELVPALGADNTLALVTQAQNLGGVDFVLIHPEFLRDRSSTSILSRLRGAGQRVAAFGWAQLGPMRDLIESANLDGWLEGPSFGQGILPAHIVALVAKMQQMRKMQQMQQIGMMTGGMNGMGGGMGGGMNGMGGGNGMNNMGNGMGGGMGNGMGGNGMNGMGGGNGMNNMGGNGMAGNGMGGGMGGNGMGGSMNGMSSGVVANVTPSAAGGMGGMMNGGMAAPQSPGMNGGRLGTNPLFNAAPSPLSSQLGAEAGMGSMGGMGGMSGMGGMGGMGGMGGAGAATTQAAGGNAEAEMLQNLMNEINRLKRELGE

**> 1XIO_*Anabaena* Sensory Rhodopsin [*Anabaena*]**

MNLESLLHWIYVAGMTIGALHFWSLSRNPRGVPQYEYLVAMFIPIWSGLAYMAMAIDQGKVEAAGQIAHYARYIDWMVTTPLLLLSLSWTAMQFIKKDWTLIGFLMSTQIVVITSGLIADLSERDWVRYLWYICGVCAFLIILWGIWNPLRAKTRTQSSELANLYDKLVTYFTVLWIGYPIVWIIGPSGFGWINQTIDTFLFCLLPFFSKVGFSFLDLHGLRNLNDSRQTTGDRFAENTLQFVENITLFANSRRQQSRRRV

**>1KGB_1 BR Bacteriorhodopsin [*Halobacterium salinarum*]**

QAQITGRPEWIWLALGTALMGLGTLYFLVKGMGVSDPDAKKFYAITTLVPAIAFTMYLSMLLGYGLTMVPFGGEQNPIYWARYADWLFTTPLLLLDLALLVDADQGTILALVGADGIMIGTGLVGALTKVYSYRFVWWAISTAAMLYILYVLFFGFTSKAESMRPEVASTFKVLRNVTVVLWSAYPVVWLIGSEGAGIVPLNIETLLFMVLDVSAKVGFGLILLRSRAIFG

>**XP_008722421.1 uncharacterized protein G647_00796 [*Cladophialophora carrionii* CBS 160.54] Halorhodopsin**

MGNDVFRHNGFTNTVTTNNHITAPGSDWYWTVCAVMTVSAFGFMIHSYFKPRSQRLFHYLNATICLIAAIAYFCMGSNLGWTAIEVEWVRSSPEVRGNMRQIFYVRYINWFITTPLIVVQLLLVAGLPTPTILYTLLMTEIVVINGLVGALVKSSYKWGFFTFGAVAFFFVAFAIVWDGRAYARVLGADVMRIFNILAAWIILLWTVYPVIWGVSEGGNIIPPDSEAVSYGVLDLLTKPVFGAVLIWGLRNVDLERLGIHVNDANPRVPRAAPAPKTEKDAEAAAAANNGVTAPAAATTPETAV

**>KIW73142.1 hypothetical protein PV04_01283 [Phialophora macrospora]**

MGNDVFRHNGFTNTVTTNNHITAPGSDWYWTVCSVMTVSAFGFMIHSYFKPRSQRLFHYLNATACLVAAVAYFCMGSNLGWTAIEVEWVRSWAKVHGNMRQIFYVRYIDWFITTPLIVSQLLLVAGLPTPTILYTLVMTEVVVINGLVGALVKSSYKWGFFTFGAVAFFFVAFAIVWDGRAYARVLGADILRIFNILAAWIVLLWTVYPIIWGVSEGGNIIPPDSEAVSYGILDLLTKPVFGAVLIWGLRNVDLERLGINITDVNPRTPRAAPAGKTEKDVEAAAAGNGVTAPAADATPETAV

**>XP_007752981.1 uncharacterized protein A1O7_00750 [Cladophialophora yegresii CBS 114405]**

MGNDVFRHNGFTNTVTTNNHITVRGSDWYWTVCAVMTVSAFGFMIHSYFKPRSQRLFHYLNATVCLIAAISYFCMGANLGWTAIQVEWMRSSAKVAGNMRQIFYVRYIDWFITTPLIVVQLLLVAGLPTPTILYTLLMTEVVVVNGLVGALVKSSYKWGFFTFGAVAFFFVAFAIAWDGRAYARVLGVDVLRIFNILAAWTILLWTVYPVIWGVSEGGNIIAPDSEAVSYGVLDLLTKPVFGAVLIWGLRNVDLARLGIHVNDAHPRIPRAEPAPRTEKDAEAAAASNGATATTPATTTATAPETAV

**>KAJ9609118.1 hypothetical protein H2200_006889 [Cladophialophora chaetospira]**

MASGFTNTVTTNNHITAAGSDWYWTVCSVMTVSAFGFMIHSFFKPRSQRLFHYLNATVCVVAAVAYFAMGSNLGWTAIEVEWLRSDAQVHGNLRQIFYVRYIDWFITTPLIVAQLLLTAGLPTPTILYTLVMTEVVVINGLVGALVKSSYKWGMFHLQFSHQSRTRRTNSDGTGFFTFGAVAFFFIAFTILYDGRAYARVLGADVLRIVNILAVWTTLLWTVYPIIWGVSEGGNIIPPDSEAVSYGILDLLTKPVFGAVLIWGLRTVDLERLGIHTIDNNPRVRGAATAPKTEKDAEAAAATNGVSAPAADPTPETAV

**>jgi|Tescy1|206752|fgenesh1_pg.7_#_159 [*Testicularia cyperi*] (Rh-MCM)**

MEAISDFAKRAGNEALSVNRPVADIDITTAGSSFLWAVFSVMAATGLGTMVWSLKVSRGERAFHYLSAAILATASVAYFAMASDLGATPVLVEFYNYAGDAAGGARPTRSIWYARYIDWTITTPLLLLEILLVSGLPLSTVFITIFFDLVMIITGLIGALVESTYKWGFYTFGCVAMFYVFYILYVPGLKSASHLGDDFKKAYLYSAMILTGLWFLYPIAWGLADGGNVISPNGEMVFYGVLDLLAKPGFALFHLFSLRRCNYSSLHLKSGKFSDYEDLGAAHYRNMRDGKAAEAGLAGDHHTTNVNGTTMGTGSTIEPAPAMRQAQVTHSLFRPQPCLCTTEKLSRGYLFPPYSSRSYILTIWHWIVPEMRTGGGMIRSIGLALATSPKRIIRRSFATSLRTARMVPDTEFTSALAKPLETVQSIYAKAQDKSAASVVKVVESHFRDTFSSDDARAKIPTLSPSSIASLLAPPSSNTAPSASHPLVRFRCMVQDTGLGTEVFLASHTQDGVEHTGLFGGEARFPVPSADESNGQQAFSNDNLTERTIMYAVSTPGQTEWAARAHRDRSGGPSHSSRSPASSSSTSSDGDISAQLANLSLMEDPPAPQKVQEVRERIRRAVDKGKSLGSTGKEPRLVAISKLHPPSAILAAHRKAGQLHFGENYVQEMVDKAKVLPREIRWHFVGGLQSNKGKLLASIPNLYLLETLDSIKAANVLQKALSSPDAAKRDEPLQVYLQVNTSGEDAKSGLPPITSADDDGKQSALLDLAVHVITKCPNLRFRGVMTIGAATNSANVQGEALEPKSVDDVVKANPDFERLIQTRRNLVRLLRSDDRIKASNESQVKEAYTELLDGSDSSADGGLELSMGMSADVDVAIMAGSDNVRVGTDCFGRRPGTRDEAMTGMKRELEIGPEQALVELRNQVQGAQQSTTSSNSAAIHDDGQPTQTASSRSAYPQKSPVPESNHIGALVKFNDLEAAESFKTAELIDVVGILDTGSLPQIEWQDTGAGQQGSSSEAPQVPCVHAILANSVDLNDVPAAWQAGSASFTSALSSKETRAELIEYIAGALAGDNLAAELVLLSITARIHARRAGLCLGALSLNISNFPAPPSSQASSSSSSSSSGGSVPETELYRRLAQLLPALVDIPMDLGSLNDPSKSLFPRSSGEGVGLEAGRLQLPSGTTIVINEGGMREGQLQDAGIRNIRALSFVLESHKLPYAFPYSEFEFDTDLNAVILSQGKSFLPFDIQCPLQAQSSDVQNGLYSPSSSAQSTAVPEDKLAQWRRSLLEARSLKTAQTFQIPESVSEHIQKEFVEDRRRQQAASSAAVAESHGGGGKSDAASGQEDLLRRMALVRLLAISRAQSSLTVETWNAAVELDTRLNERIQAQNQNQQSAASQPSISR
